# Plug-and-play protein biosensors using aptamer-regulated in vitro transcription

**DOI:** 10.1101/2023.08.10.552680

**Authors:** Heonjoon Lee, Tian Xie, Xinjie Yu, Samuel W. Schaffter, Rebecca Schulman

## Abstract

Molecular biosensors that accurately measure protein concentrations without external equipment are critical for solving numerous problems in diagnostics and therapeutics. Modularly transducing the binding of protein antibodies, protein switches or aptamers into a useful output remains challenging. Here, we develop a biosensing platform based on aptamer-regulated transcription in which aptamers integrated into transcription templates serve as inputs to molecular circuits that can be programmed to a produce a variety of responses. We modularly design molecular biosensors using this platform by swapping aptamer domains for specific proteins and downstream domains that encode different RNA transcripts. By coupling aptamer-regulated transcription with diverse transduction circuits, we rapidly construct analog protein biosensors or digital protein biosensors with detection ranges that can be tuned over two orders of magnitude. Aptamer-regulated transcription is a straightforward and inexpensive approach for constructing programmable protein biosensors suitable for diverse research and diagnostic applications.

**One sentence summary:** We develop a modular platform for biosensing across a wide dynamic range using aptamer-regulated transcription to detect different proteins and molecular circuits to process the RNA transcript outputs.

## Introduction

Biosensors that detect proteins are crucial for diagnostics^1^, smart therapeutics^2^, and biomedical research. Traditional ways to detect or measure the concentrations of proteins, such as enzyme-linked immunosorbent assay (ELISA)^3^ or Western blot^4^ often rely on specific and sensitive binding of antibodies to a target protein. However, these assays are time-consuming and labor-intensive due to multiple incubation and washing steps. Mass spectrometry^5^, single-molecule microscopy^6^, and electrochemical sensors^7^ can detect diverse proteins, but these analytics require expensive and bulky instrumentation, infrastructure, and trained personnel. Further, the output signals of such assays cannot be easily tailored to applications besides protein quantification such as portable diagnostics^8,9^ or molecular machines that perform targeted delivery^10,11^ in response disease biomarker detection. Lateral flow assays^12^ that rely on antibody binding permit low-cost and rapid detection of proteins, but typically only report the presence of a biomarker and cannot be integrated with downstream reactions that process the input signals^13,14^.

Ideally, molecular biosensors could generate a response to a target protein without manual intervention, could be read without specialized equipment, and could easily be programmed to produce a variety of responses. Antibodies^15^, protein switches^16^, and aptamers^17,18^ can be designed to bind to specific proteins, but modularly designing circuits that transduce the protein binding event into a measurable output is challenging. For example, allosteric protein switches^19,20^ or aptamers^21,22^ that undergo conformational changes upon protein binding to transduce signals have been integrated with molecular circuits. However, designing these structure-switching molecules often requires extensive reengineering for each target protein or desired output^23^. This makes it difficult to rapidly develop protein biosensors against new targets or to adopt existing biosensors for new functionalities.

Here, we present <u>a</u>ptamer-<u>r</u>egulated transcription for *in vitro* <u>s</u>ensing and transduction (ARTIST), which enables rapid construction of protein biosensors that can detect diverse targets and easily integrate with downstream circuits for programmable responses. ARTIST uses aptamer-protein binding to regulate transcription of a DNA template in analogy to the regulation of transcription by protein transcription factor-DNA binding^9,24^ (Figure 1a). Using published DNA aptamer sequences, we show how to simply swap the aptamer domain in these DNA templates for aptamer-regulated transcription (dARTs) to selectively sense a range of proteins, including multiple cytokines involved in autoimmune diseases. Conversely, we modularly swap dART output domains to integrate dARTs with different molecular circuits to create biosensors with different capabilities (Figure 1b).

**Figure 1.**
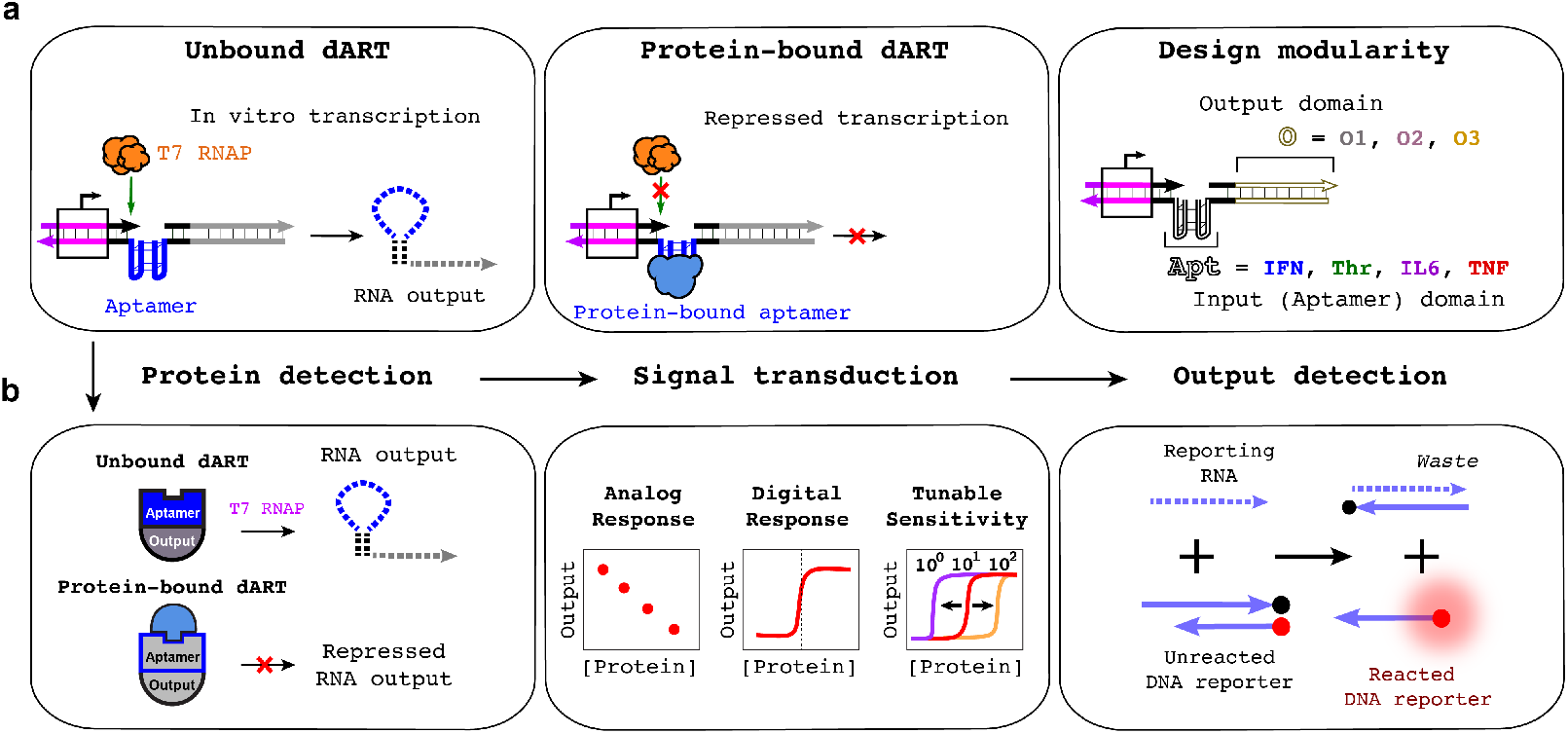
The ARTIST platform. **(a)** In the absence of an aptamer’s protein ligand, dARTs are transcribed to produce an RNA output (left), but protein binding represses transcription (middle). The input and output domains are decoupled (right), which enables modular design of dARTs by swapping out the aptamer domain or customizing the output sequences to encode different RNA outputs. DNA and RNA are represented with solid and dashed lines, respectively. IFN, Thr, IL6, and TNF indicate DNA aptamers for IFN-γ, thrombin, IL-6, and TNF-α, respectively. **(b)** dARTs serve as the protein sensing layer (left) whose outputs can be coupled with downstream circuits to demonstrate versatile functionalities (middle). The RNA output of molecular circuits can react with a DNA reporter complex (right), which produces measurable fluorescence for detection.

Using ARTIST, we build analog biosensors whose outputs indicate protein concentration and digital biosensors in which substantial output is only produced when the protein’s concentration exceeds a chosen threshold. The dynamic range of these biosensors can be rationally tuned across multiple orders of magnitude to detect cytokines at physiologically relevant concentrations; even below the aptamer *K*_*d*_, which often limits biosensor sensitivity in practice. We also demonstrate how ARTIST can be used to create biosensors that are robust to environmental parameters that biosensors are often sensitive to^25^, such as ion concentration and enzyme activity. The ARTIST platform should enable the facile development of rapid (<60 min) point-of-care diagnostics involving diverse proteins.

## Results

### dART design and characterization

The formation of noncanonical DNA structures such as G-quadruplexes have been shown to regulate transcription in cells^26–28^ and have been adapted to inhibit transcription *in vitro*^29,30^. We hypothesized we could develop a general protein biosensing platform by designing DNA transcription templates, termed dARTs, with G-quadruplex-forming aptamers downstream of a promoter that repress transcription *via* protein-aptamer binding. dARTs consist of a promoter domain that T7 RNA polymerase (T7 RNAP) can recognize to initiate transcription, a single-stranded aptamer domain that can bind to a specific protein, and an output domain that T7 RNAP transcribes to produce an RNA output sequence that can react downstream (Figure 2a). The aptamer sequence was inserted into the template strand of the dART ^29,30^, *i*.*e*. the strand the polymerase reads during transcription. The output domain of the dART was designed to be double-stranded to prevent spurious interactions between the dART and other species such as the RNA transcribed from the dART. The aptamer and the output sequences are both transcribed from dARTs, so the resulting transcripts could have undesired secondary structure (Figure S1, Supplementary Information Section 1.1). To reduce the possibility of such interactions, we added insulation domains, composed of *i* and *i’*, to the dART design (Figure 2b). The *i* and *i’* domains are complementary and are located on either side of the aptamer sequence so that in the RNA transcript produced from the dART, these domains will hybridize to form a hairpin, sequestering the sequence transcribed from the aptamer domain. (Figure 2a, Supplementary Information Section 1.2)^31,32^. The *i* domain located just downstream of the promoter was designed to be 6 bases to facilitate efficient T7 RNAP transcription initiation^33^.

**Figure 2.**
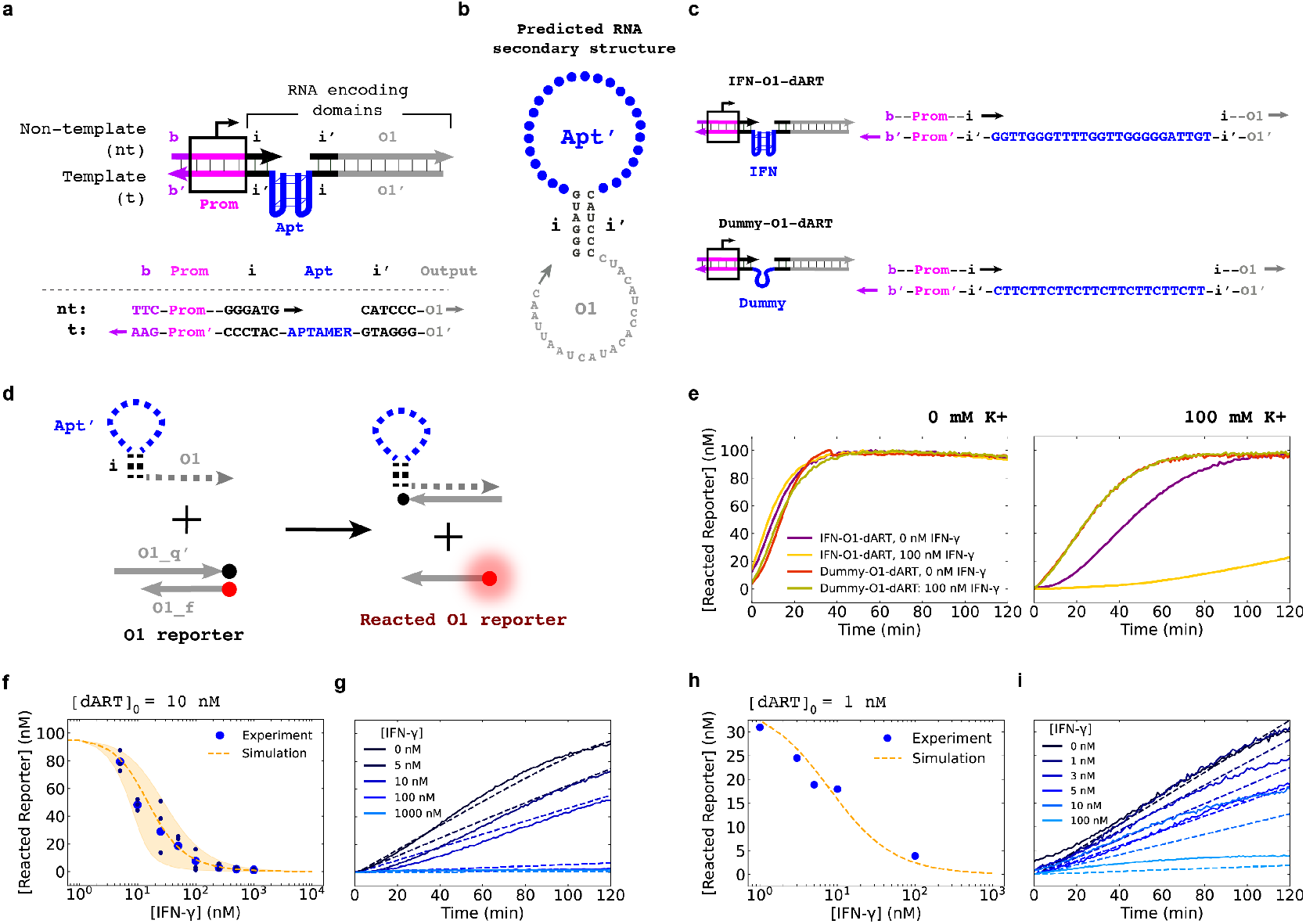
Design of aptamer regulated transcription templates (dARTs). **(a)** dART templates consist of a double-stranded promoter region (pink boxed region), a single-stranded aptamer domain on the template strand read by T7 RNAP (Apt), and a double-stranded output domain (O1). The single-stranded aptamer domain permits aptamer-ligand binding. (**b**) Secondary structure of the RNA transcript predicted by NUPACK^38^. **(c)** The aptamer sequences of IFN-O1-dART, which binds IFN-γ and Dummy-O1-dART, whose aptamer domain does not have a tertiary structure or specific protein affinity. **(d)** dART transcripts react with an O1 DNA reporter via toehold-mediated strand displacement. **(e)** Reacted reporter kinetics from IFN-O1-dART and Dummy-O1-dART transcription with and without IFN-γ and potassium. **(f)** Simulated and experimental dose-response curves for 10 nM IFN-O1-dART with 0 nM to 1000 nM IFN-γ for *K*_*d, apparent*_ = 8 nM (Supplementary Information Section 2). Three independent replicates (black) and their mean (blue) are plotted. The dashed line is a simulation for *K*_*d, apparent*_ matching experiments. The shaded region shows simulation results for K_d, apparent_ values spanning 1 nM to 20 nM (see Methods). **(g)** Experimentally measured (bold) and simulated (dashed) reacted reporter kinetics for 10 nM of IFN-O1-dART and 0 nM to 1000 nM of IFN-γ. **(h)** Simulated and experimental dose-response curves and **(i)** Experimental (bold) and simulated (dashed) reacted reporter kinetics **(h)** for 1 nM IFN-O1-dART and 0 nM to 100 nM of IFN-γ using the *K*_*d,apparent*_ determined in **(f)**. For **(f)** and **(h)**, the reacted reporter concentration for each plot was measured at 120 min. See Supplementary Information Section 2 for model equations and parameters considered to determine *K*_*d,apparent*_.

We next sought to characterize how our dART design performed in different experimental conditions. For a protein to reduce the rate of dART transcription, the protein must be able to bind to the dART’s aptamer and repress transcription, while under the same conditions dARTs should be efficiently transcribed when no protein is present. We began by designing a dART, termed the IFN-O1-dART, that contained an IFN-γ aptamer that forms a G-quadruplex to facilitate IFN-γ binding (Figure 2c)^34,35^. For characterization, we used a fluorescently labeled DNA reporter complex that reacts with IFN-O1-RNA *via* toehold-mediated strand displacement (TMSD) to displace one strand of the reporter complex to produce a fluorescent signal (Figure 2d, Methods, Figure S2, Supplementary Information Section 1.3).

Many G-quadruplex aptamers require potassium ions for effective ligand binding^36,37^. Consistent with this requirement, in the absence of K+, the same rate of fluorescence decrease (*i*.*e*. the rate of transcript reaction with the reporter) was observed for IFN-O1-dART in either the presence of 100 nM of IFN-γ or its absence (Figure 2e), suggesting IFN-γ did not bind to the IFN-O1-dART to reduce transcription rate. When 100 mM KCl was also added, 100 nM IFN-γ was able to substantially repress transcription, consistent with IFN-γ binding to the dART. Without IFN-γ present, the rate of reporter reaction of the IFN-O1-dART was slightly lower with 100 mM of potassium than without potassium, which is consistent with the ability of G-quadruplexes to interrupt transcription^26,30^. As a control, we verified that a dART with a “dummy” aptamer domain (Dummy-O1-dART), i.e., no specific affinity for IFN-γ and no G-quadruplex forming domain, had similar reacted reporter kinetics for 0 nM or 100 nM of IFN-γ at both 0 mM and 100 mM of K+ (Figure 2c,e). We also tested dART variants with *i* domain lengths ranging from 2 bases to 22 bases (Figure S3, S4, Supplementary Information Section 1.4) and found the 6 base design yielded the greatest difference in transcription rates with and without protein in our experimental conditions (Figure S5).

We next developed a simple model to predict the dose-response and kinetic behavior for 10 nM of IFN-O1-dART incubated with IFN-γ concentrations ranging between 0 nM and 1000 nM. Our model assumes that transcription cannot occur when a protein is bound to a dART’s aptamer, and thus the apparent dissociation constant of the protein and the dART’s aptamer, which we termed *K*_*d,apparent*_, will dictate the concentration of dART available for transcription at a given protein concentration (Supplementary Information Section 2). This model qualitatively agrees with our measured dose-response curve for a *K*_*d, apparent*_ of 8 nM (Figure 2f and Figures S6-S8), which is on the same order as the measured *K*_*d*_ of the dART’s aptamer sequence^34,35^. Simulations of reacted reporter kinetics were also in good agreement with those measured in experiment (Figure 2g). Using the model, we then compared the predicted dose-response curve using the *K*_*d,apparent*_ of 8 nM to the experimental dose-response curve of 1 nM of IFN-O1-dART in the presence of (0, 1, 3, 5, 10, or 100) nM of IFN-γ. Simulated and experimental results were consistent with each other for both dose-response curve (Figure 2h) and dose-dependent kinetic profiles (Figure 2i).

### dART input and output modularity

We next asked whether the design of the IFN-O1-dART could be generalized to respond to different protein inputs and produce different RNA outputs. We replaced the IFN-γ aptamer in the IFN-O1-dART with G-quadraplex forming aptamers for thrombin^39^, IL-6^40^, and TNF-α^41^ to create Thr-O1-dART, IL6-O1-dART, and TNF-O1-dART (Figure 3a, Figure S9, Supplementary Information Section 3.1). NUPACK^38^ predicted that the insulation domains would prevent undesired secondary structures from forming in these new output RNA sequences (Figure S10).

**Figure 3:**
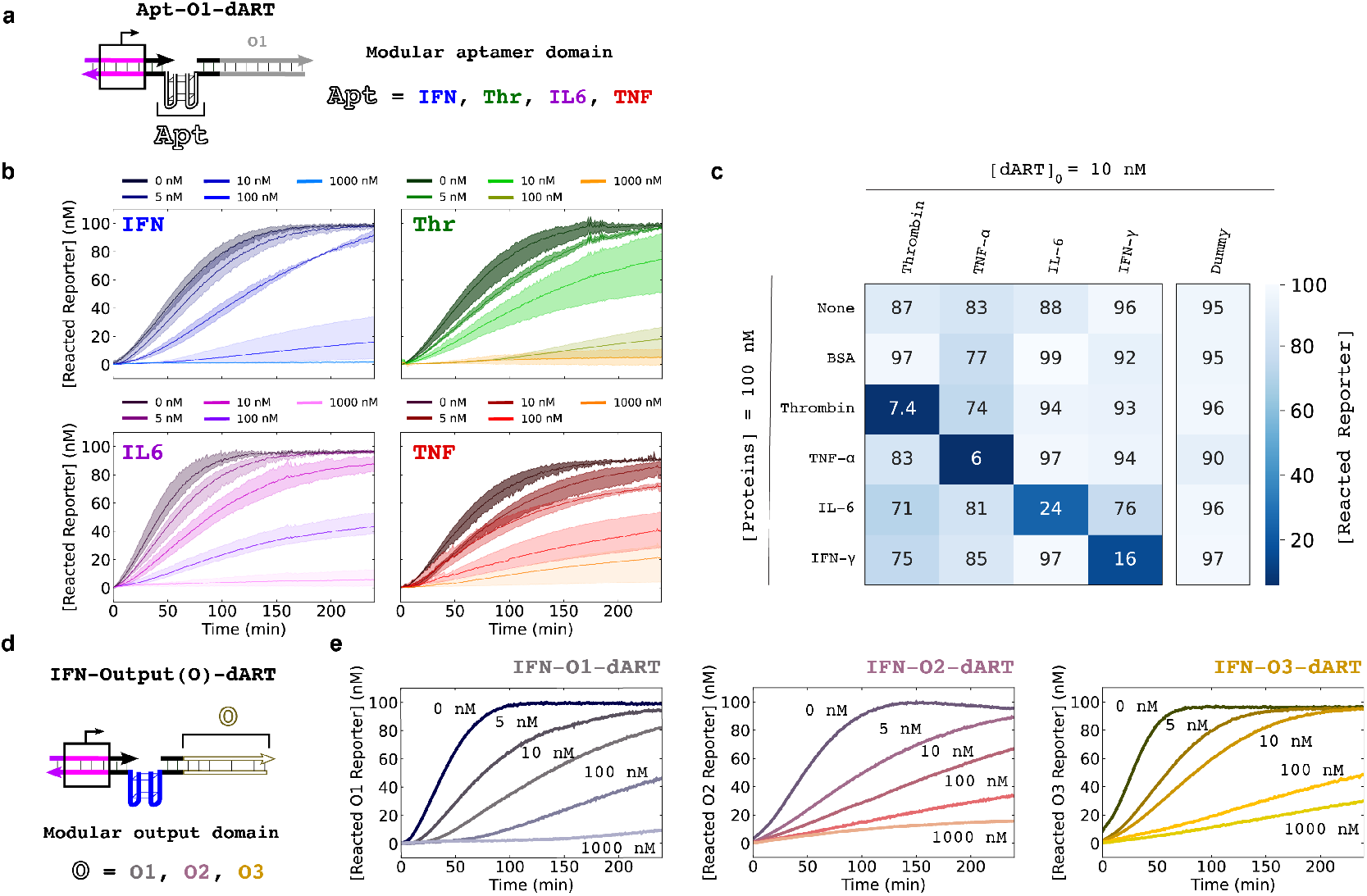
dART sensitivity, selectivity, and modularity. **(a)** A series of dARTs with different inputs, achieved by swapping aptamer domain sequences. **(b)** Reacted reporter kinetics by IFN-O1-dART, Thr-O1-dART, IL6-O1-dART, and TNF-O1-dART in the presence of 0 nM to 1000 nM IFN-γ, thrombin, IL-6, and TNF-α, respectively. Shaded regions represent minimum/maximum values of reacted reporter concentration of three independent replicates. **(c)** Heat map showing reacted reporter concentrations for 10 nM Thr-O1-dART, IFN-O1-dART, IL6-O1-dART, TNF-O1-dART, and Dummy-O1-dART (a control) each subjected to 100 nM of BSA, thrombin, TNF-α, IL-6, or IFN-γ. **(d)** A series of dARTs with different outputs, achieved by swapping template output sequences. **(e)** Reacted reporter kinetics for IFN-O1-dART, IFN-O2-dART, and IFN-O3-dART for 0 nM to 1000 nM of IFN-γ. All experiments were conducted in the presence of 100 mM KCl. Sequences of IFN-O1-dART, Thr-O1-dART, IL6-O1-dART, TNF-O1-dART, IFN-O1-dART, IFN-O2-dART, and IFN-O3-dART are in Supplementary Information Section 3.

We measured the reacted reporter kinetics for 10 nM of each of these dARTs when combined with 0 nM to 1000 nM of their target proteins (Figure 3b). For all dARTs, the rates of reacted reporter decreased with increasing protein concentrations. We next asked whether these changes in dART reporter reacted were due to specific binding between a target protein and its aptamer. We combined 10 nM of each dART and Dummy-O1-dART with 100 nM of each protein ligand and BSA as a control. Only when dARTs were subjected to their target protein was there a large decrease in the reacted reporter concentration (Figure 3c and Figure S11). As expected, the dummy template produced a similar concentration of reacted reporter in the presence of all input proteins.

We next asked whether the output domain could also be exchanged modularly, which would allow dARTs to be easily connected to different downstream processes (Figure 3d, Supplementary Information Section 3.2). We measured transcription profiles of IFN-O1-dART, IFN-O2-dART, and IFN-O3-dART which encode the O1, O2, and O3 output domains, respectively, for 0 nM to 1000 nM of IFN-γ (Figure S12 and S13). dARTs with each output resulted in the expected reacted reporter dose-response curve across IFN-γ input concentrations (Figure 3e), suggesting that the outputs of the dARTs could be easily customized to couple them to downstream circuits to create biosensors with different functionalities.

Not all aptamer domains integrated well with dARTs. For example, when we introduced aptamers for kanamycin^42^ and VEGF^43^ (Supplementary Information Section 3.3, Figures S14), we observed negligible reacted reporter even in the absence of their corresponding ligands (Figure S15). This could be because the aptamers form stable enough G-quadruplexes with K+ to repress transcription even without ligand, or because the RNA outputs form undesired secondary structure that impede their reaction with the reporter (Figure S16).

### Analog biosensors

We next asked whether we could use ARTIST to build biosensors with different functionalities, such as analog or digital response, or signal amplification by coupling dARTs to downstream reactions. In principle, the rate of increase of reacted reporter by a dART is a measure of the target protein’s concentration. However, rates are difficult to measure because they require observing a signal over time. We thus asked whether we might build analog biosensors that indicate protein concentration as the steady-state concentration of reacted reporter by adding RNase H. RNase H degrades RNA outputs of a dART bound to the DNA strand O1’_q^31,44^, allowing O1’_q to rehybridize to O1_f to reform the quenched reporter complex after degradation (Figure 4a). A balance of RNA production and degradation should therefore produce a steady-state response whose magnitude is dependent on the concentration of the target protein. The reacted reporter concentrations of IFN-O1-dART with 0 nM to 1000 nM of IFN-γ indeed reached different steady-state values for different input protein concentrations in the presence of RNase H (Figure 4b-c). When Thr-O1-dART, IL6-O1-dART, and TNF-O1-dART were combined with reporter and 5 nM to 1000 nM of their corresponding input proteins, distinct steady-state reacted reporter concentrations were also observed (Supplementary Information Section 4, Figure S17, Figure 4c).

**Figure 4.**
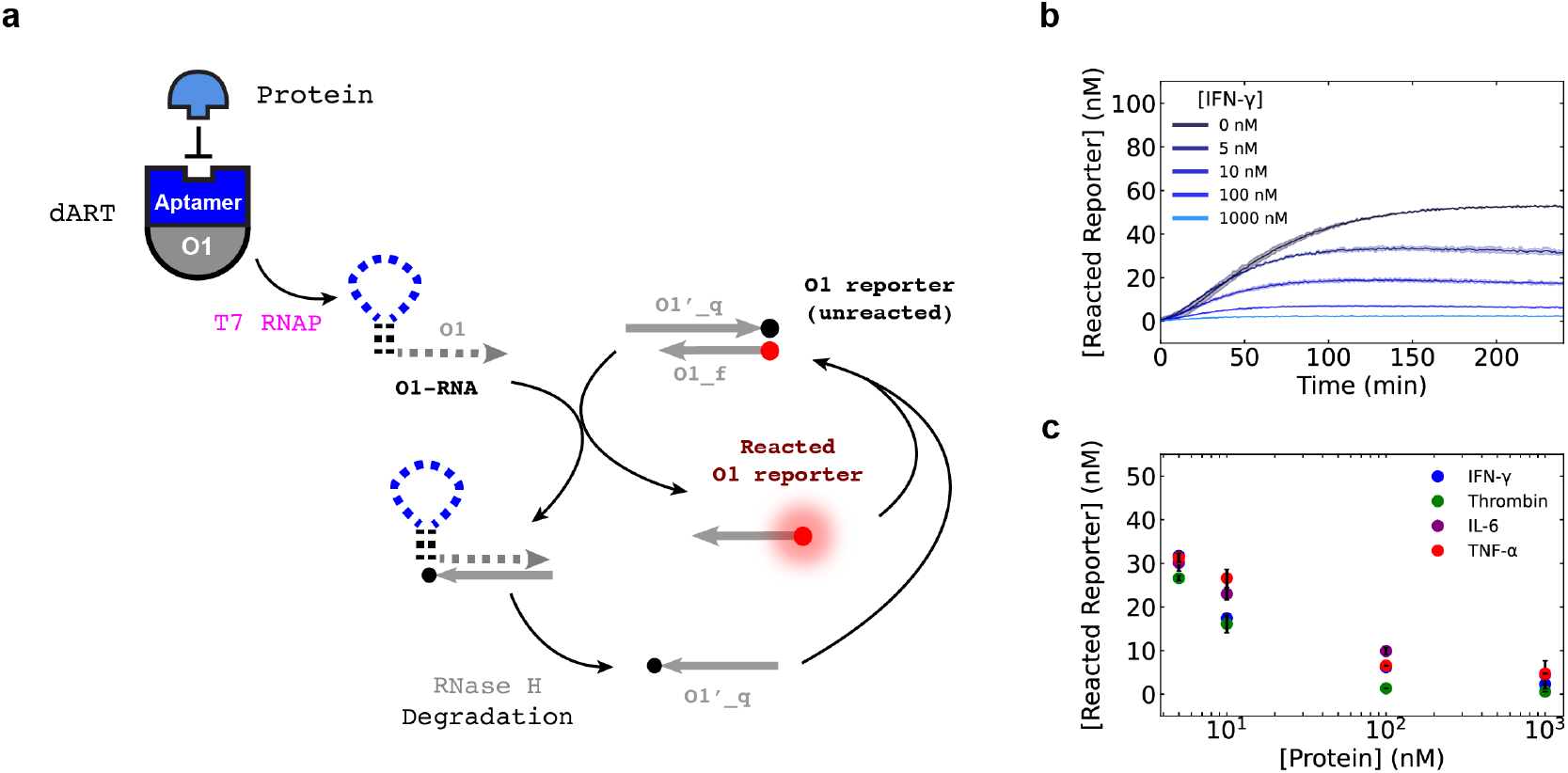
Analog biosensors with steady-state outputs. **(a)** Schematic of coupled RNA transcription of dARTs by T7 RNAP and RNA degradation by RNase H. **(b)** The concentration of reacted reporter over time for an experiment with 10 nM IFN-O1-dART, 0 nM to 1000 nM of IFN-γ, 2 U μL^-1^ of T7 RNAP, and 2 × 10^−3^ U μL^-1^ of RNase H. Shaded regions enclose the minimum and maximum values for independent trials (N=2; see Methods). **(c)** Steady-state analog outputs of IFN-O1-dART, Thr-O1-dART, IL6-O1-dART, and TNF-O1-dART with 5 nM to 1000 nM of their corresponding proteins. Error bars represent minimum and maximum values of two independent replicates.

### Digital biosensors

We next asked how we might use ARTIST to construct digital biosensors, which produce a high output signal when the input exceeds a set threshold concentration and a low output signal otherwise. Digital biosensors are important for measuring whether a sample satisfies a specific diagnostic criterion. We designed a digital biosensor for IFN-γ by integrating two dARTs to produce a comparator circuit. A reference dART with an aptamer domain that does not bind IFN-γ was designed to produce the O1 output (Ref-O1-dART). Another dART with the IFN-γ aptamer domain was designed to produce an output with partial complementarity to the O1 sequence (IFN-O1’-dART) (Figure 5a, Supplementary Information Section 5.1, Figure S18). If the IFN-O1’-dART is added at a higher concentration than the Ref-O1-dART, then IFN-O1’-RNA will sequester Ref-O1-RNA, resulting in a low reporter signal in the absence of protein. As IFN-γ concentration is increased, the rate of IFN-O1’-RNA transcription will decrease, allowing the Ref-O1-RNA to react with the O1 reporter once a threshold IFN-γ concentration is reached (Figure 5b). This comparator circuit also inverts the output signal of individual dARTs, so that high protein concentration yields high output signal.

**Figure 5.**
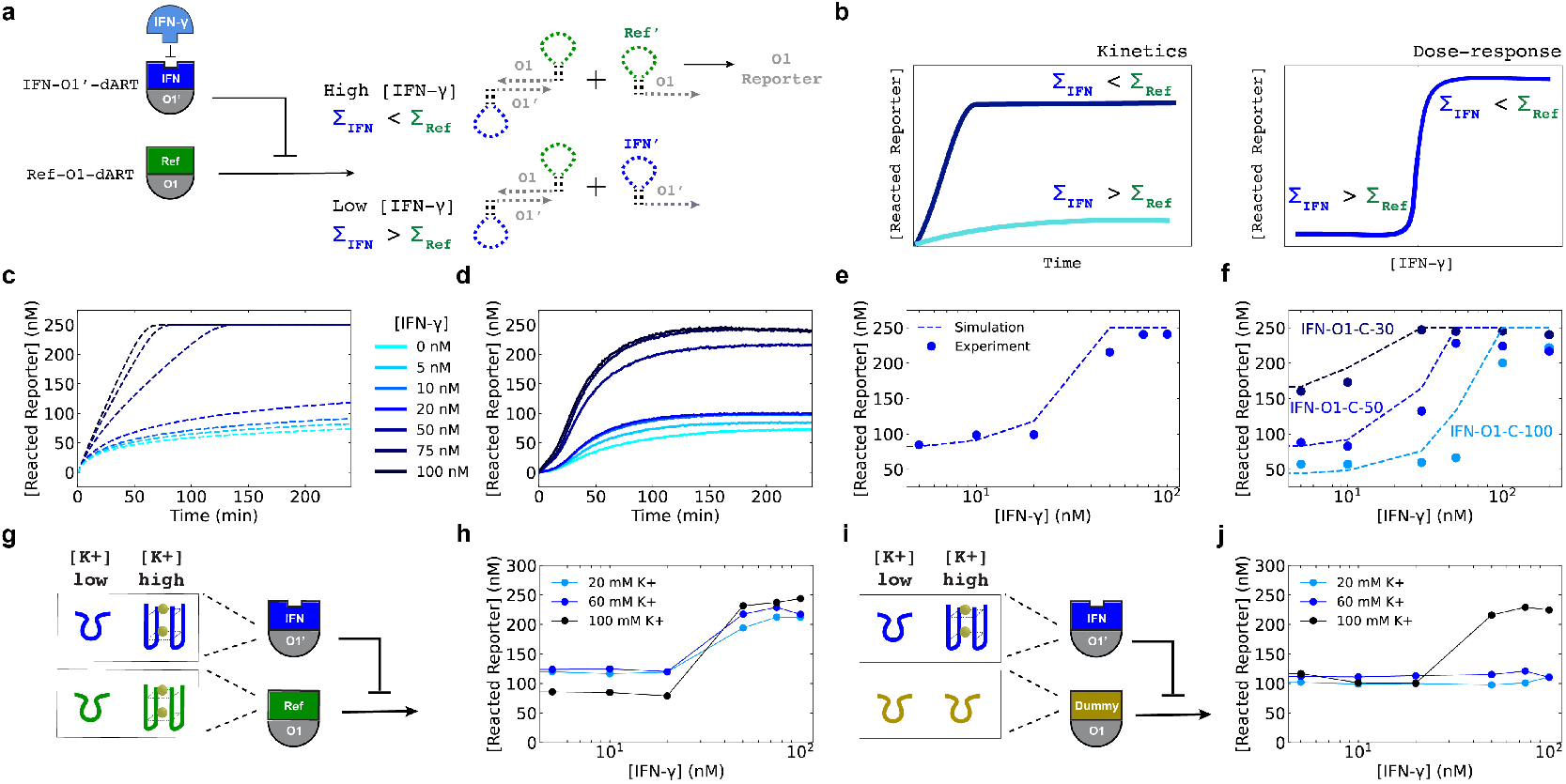
Comparator circuits for digital biosensing. **(a)** The comparator circuit. When [IFN-γ] is high, IFN-O1’-dART’s transcription rate is low. Ref-O1-RNA is thus produced in excess, and it reacts with the O1 DNA reporter. When [IFN-γ] is low, IFN-O1’-RNA’s transcription rate is higher than Ref-O1-RNA’s, and Ref-O1-RNA is sequestered by IFN-O1’-RNA before it can react with the reporter. **(b)**. Illustrations of desired responses of the digital biosensor. Left: when [IFN-γ] is high, reacted reporter rises concentration should increase rapidly until all reporter is reacted, When [IFN-γ] is low it should rise slowly or not at all. Right: the concentration of reacted reporter should therefore either be fully ON or very low (OFF) at the end of the reaction. **(c)** Simulated kinetics of IFN-O1-C-50. **(d)** Experimentally measured kinetics of IFN-O1-C-50. **(e)** Dose-response curve of IFN-O1-C-50 after 240 min of reaction. Dashed lines represent simulations, whereas the points represent experimental values. **(f)** The dose-response curves of IFN-O1-C-30, IFN-O1-C-50, and IFN-O1-C-100. **(g)** The comparator is designed to be K+-insensitive. IFN-O1’-dART and Ref-O1-dART both form G-quadruplexes, and their transcription rates should change similarly with [K+] concentrations so the output of the comparator should be relatively insensitive to [K+]. **(h)** Dose-response curves of IFN-O1-C-50 in (g); responses were measured after 240 min. The line plots are added to aid visualization of the trends in the reacted reporter concentrations for each [K+] concentration. **(i)** A [K+]-sensitive comparator. At high [K+], the aptamer domain on the IFN-O1’-dART forms a G-quadruplex, which reduces its transcription rate. If the IFN-O1’-dART’s output is compared to an RNA produced at a rate insensitive to [K+] (from Dummy-O1-dART) output should change with [K+]. **(j)** Dose-response curves of IFN-O1-C-50 with Dummy-O1-dART for different [K+]; responses were measured after 240 min. The line plots are added to aid visualization of the trends for each [K+] concentration. See Supplementary Information Sections 5.2, 5.3, and 5.4 for simulation equations and parameters.

We first sought to build a comparator circuit that could threshold the concentration of IFN-γ so that [IFN-γ] < 20 nM would produce a low reacted reporter signal (OFF) and [IFN-γ] ≥ 50 nM would produce a high signal (ON). We used simulations with the previously measured *K*_*d,apparent*_ for the IFN-O1-dART (Supplementary Information Sections 5.2, 5.3, and 5.4, Figure S19) to determine that 25 nM of Ref-O1-dART and 50 nM of IFN-O1’-dART should produce a digital response with the desired threshold (Figure S20 and Figure 5c). These predictions were then confirmed in experiments (Figure 5d,e). We termed this IFN-γ O1 comparator circuit involving 25 nM of Ref-O1-dART and 50 nM of IFN-O1’-dART, IFN-O1-C-50.

We reasoned we could tune the IFN-γ threshold of IFN-O1-C-50 by changing the concentration of Ref-O1-dART, which would change the amount of IFN-O1’-dART that must be repressed by input protein to produce a reporter signal (Supplementary Information Section 5.4)^13^. In line with this intuition, we found that an IFN-γ O1 comparator circuit with 15 nM of Ref-O1-dART required ≥ 100 nM IFN-γ to turn on (IFN-O1-C-100), while a circuit with 40 nM Ref-O1-dART only required ≥ 30 nM IFN-γ to turn on (IFN-O1-C-30; Figure 5f). In principle, these circuits could measure the activation of CD4 T-cells, which can secrete above 30 nM of IFN-γ within microwells after 15 hours of mitogenic stimulation^45^.

### Robust digital responses

We hypothesized that the digital response of IFN-O1-C-50 would also be insensitive to different environmental factors that often affect biosensors. For example, T7 RNAP activity can vary with solution composition^46^ or temperature^47^. Since a comparator computes the ratio of transcription rates, the digital output should not change substantially with T7 RNAP activity, which would change both transcription rates. Indeed, the IFN-O1-C-50 with T7 RNAP activities of 2 U μl^−1^, 4 U μl^−1^, and 8 U μl^−1^ all produced low reacted reporter concentrations for [IFN-γ] < 20 nM and high reacted reporter concentrations for [IFN-γ] ≥ 50 nM (Figures S21 and S22, Supplementary Information Section 5.5).

The affinity of G-quadruplex-forming aptamers for their ligands is also sensitive to potassium ion concentrations^36,48^, and we found that dART transcription rate decreased with increasing potassium concentration (Supplementary Information Section 5.6, Figure S23). Despite the potassium sensitivity of individual dARTs, both dARTs comprising IFN-O1-C-50 contain G-quadraplexes (Figure 5g), so their transcription rates are affected in similar ways by different potassium concentrations, resulting in similar digital responses and response thresholds for IFN-O1-C-50 for different potassium concentrations (Figure 5h). In contrast, a comparator with a reference dART with a dummy aptamer sequence (Figure 2c, Supplementary Table 1) that does not form a G-quadruplex (Figure 5i), becomes non-responsive at lower potassium ions concentrations (Figure 5j, Figure S24) because the IFN-O1’-dART transcription rate changes with potassium concentration, but the Dummy-O1-dART transcription rate does not, changing the threshold. These results show how ARTIST can be used to construct digital biosensors that maintain their desired response profiles under different environmental conditions, which may eventually make it possible to construct sensors useful for different biological samples such as blood, urine, or saliva.

### Output amplification

We next asked whether we might increase the amount of output produced by a comparator circuit in response to the same concentrations of protein input. We designed an amplified comparator (Figure 6a, Supplementary Information Section 6) in which the RNA output of the comparator circuit activates transcription of a downstream genelet^31^ via strand displacement reactions that complete the promoter sequence of the genelet (Figure 6b, Figure S25, S26). To prevent the low level of comparator output that is produced even when [IFN-γ] is low from triggering the output, we included RNase H, so that only when the rate of Ref-C1-RNA production exceeds its degradation rate can it activate the genelet to produce an output.

**Figure 6.**
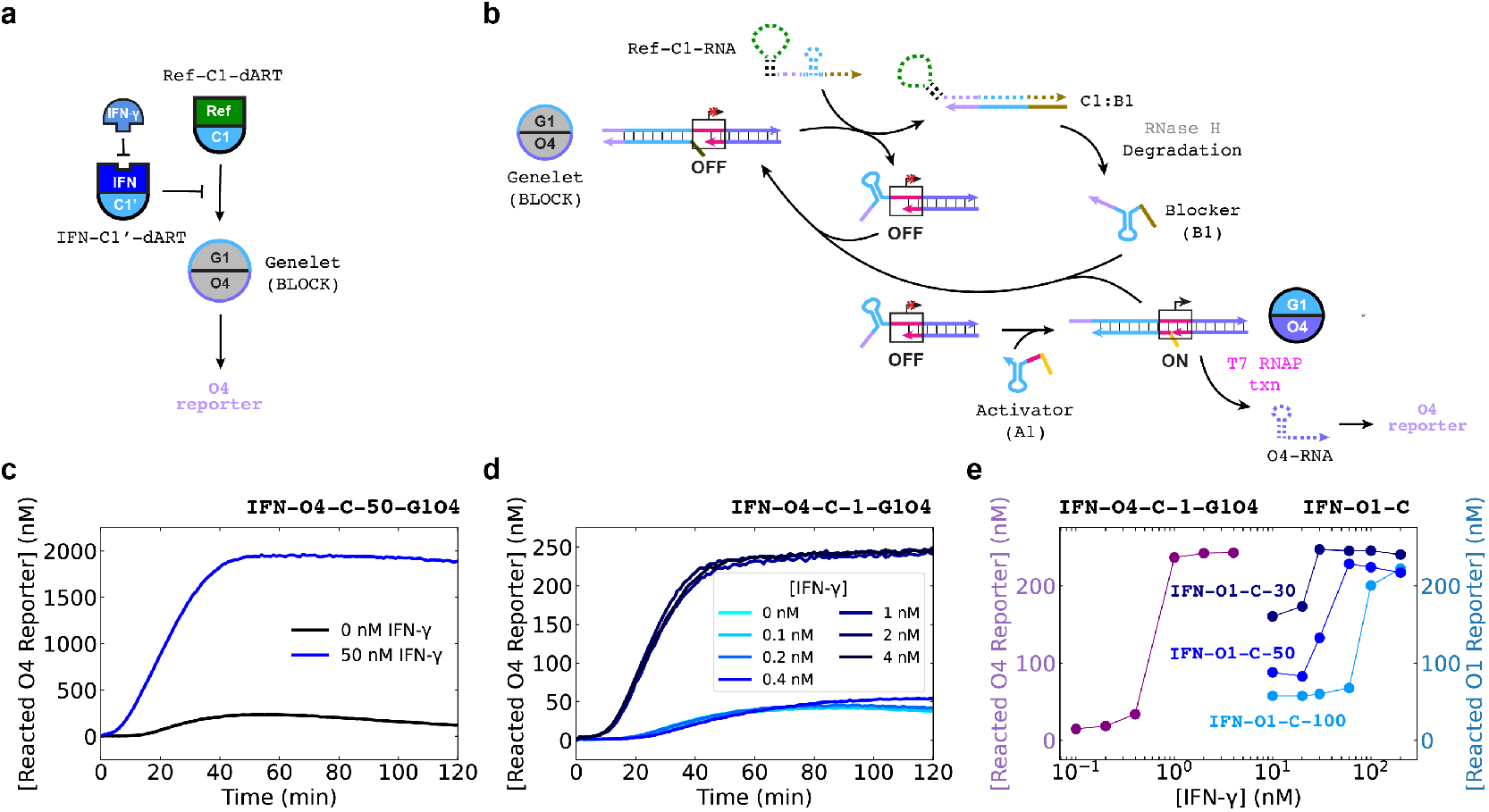
Sensitive protein detection with amplified comparators. **(a)** Circuit diagram for an amplified IFN-C1’-dART / Ref-C1-dART comparator. The Ref-C1-RNA output, which is high when IFN-γ exceeds a threshold concentration, activates transcription of a genelet that transcribes O4, which is detected by a reporter. **(b)** The reactions through which Ref-C1-RNA coactivates G1O4 to produce the O4 output. **(c)** Reacted reporter kinetics of IFN-O4-C-50-G1O4, which consists of 25 nM Ref-C1-dART, 50 nM IFN-C1’-dART, 100 nM G1O4:B1, 200 nM A1, and 2000 nM O4 DNA reporter, for 0 nM or 50 nM IFN-γ. **(d)** Reacted reporter kinetics of the IFN-O4-C-1-G1O4, which consists of 0.5 nM of Ref-C1-dART, 1 nM IFN-C1’-dART, 20 nM of G1O4:B1, and 100 nM of A1, and 250 nM of O4 DNA reporter for IFN-γ inputs from 0 nM to 4 nM. 4 U μL^-1^ T7 RNAP and 4 × 10^−3^ U μL^-1^ RNase H were used in IFN-O4-C-1-G1O4. **(e)** Comparison of the dose-response curves of IFN-O4-C-1-G1O4, IFN-O1-C-30, IFN-O1-C-50, and IFN-O1-C-100. The lines connecting points were added to aid visualization of the trends for each dose-response curve.

The amplified comparator (IFN-O4-C-50-G1O4) reacted with 2000 nM of O4 reporter (Figure 6c) in response to 50 nM of IFN-γ in <45 min, more than 40 times the output produced by the non-amplified comparator to the same input (IFN-O4-C-50; Figure S27). Without IFN-γ, about 200 nM of O4 reporter reacted transiently, but this amount then decreased, presumably because of RNase H-catalyzed O4 RNA degradation. Interestingly, the ON/OFF ratio of IFN-O4-C-50 was only 2-fold to 3-fold (Figure S27), compared to IFN-O4-C-50-G1O4, which was nearly 10-fold (Figure 6c). This increase in the fold-change between the ON and OFF signals may be due to two factors. First, Ref-O1-RNA reacts faster with the O1 reporter in IFN-O4-C-50 than Ref-C1-RNA does with B1 in IFN-O4-C-50-G1O4^31^. Thus, when IFN-C1’-RNA is in excess, a greater fraction of Ref-C1-RNA is sequestered in IFN-O4-C-50-G1O4 than IFN-O4-RNA in IFN-O4-C-50. Second, in IFN-O4-C-50-G1O4, both Ref-C1-RNA and O4-RNA are degraded by RNase H, which results in a lower background signal without protein for IFN-O4-C-50-G1O4 than for IFN-O4-C-50.

We hypothesized that an amplified comparator such as IFN-O4-C-50-G1O4 could be tuned to produce a similar amount of high output to its corresponding non-amplified comparator (IFN-O4-C-50) beginning above a much lower threshold protein concentration. To build this more sensitive amplified comparator, we reduced the concentrations of the Ref-C1-dART and IFN-C1’-dART of IFN-O4-C-50-G1O4 50-fold to create a diluted amplified comparator, termed IFN-O4-C-1-G1O4. IFN-O4-C-1-G1O4 maximized reporter signal for IFN-γ concentrations of 1 nM or higher and otherwise produced < 12 % of the maximum reporter signal. Amplifying the output of the comparator thus makes it possible to reduce the threshold input concentration of the circuit 50-fold (Figure S28), while producing a similar amount of output when ON (Figure 6d-e). IFN-O4-C-1-G1O4’s 1 nM responsive threshold is lower than the apparent dissociation constant for the IFN-γ dART that we measured (8 nM). Furthermore, the amount of reacted reporter, a measure of the RNA output, produced by IFN-O4-C-1-G1O4 at this threshold is >250-fold higher than the input protein concentration (Figure 6d-e).

## Discussion

ARTIST is a versatile platform for rapidly developing biosensors that report on protein concentrations. By coupling downstream reactions to dART’s protein-controlled transcription processes, ARTIST can be easily tailored to produce stable analog or digital responses, to increase signal-to-noise ratios, or to increase sensitivity by almost two orders of magnitude.

ARTIST is already applicable for sensing proteins of research and clinical interest. For example, IFN-γ is a ubiquitous cytokine involved in various malignancies (e.g., cancer, autoimmune diseases)^49^, as well as a key signaling factor in immunotherapy. Different aptamers that have been selected from previous studies can readily be integrated into dARTs, suggesting the promise of developing a larger library of biosensors for a range of important protein targets. In addition to the downstream reactions studied here, dART outputs could be processed using a broad range of available RNA amplification^50^ or sequencing^51^ methods. The selectivity of dARTs suggests that they may be combined to allow for multiplexed sensing or decision-making involving multiple protein inputs. The constituents of ARTIST, T7 RNAP, RNase H, and DNA complexes can be easily freeze-dried for storage and distribution^8,9^; these methods could provide a route to creating portable biosensors for point-of-care use.

Nonetheless we found two aptamers that could not be easily integrated into dARTs (Supplementary Information Section 3.3). It will be important to better understand the criteria that ensure the successful integration of aptamers into dARTs. Both aptamers that did not work showed low transcription in the absence of ligand. One possibility is that aptamers with G-quadruplexes that are highly stable even in the absence of ligand may inhibit dART transcription on their own^48^. For such aptamers, it might be possible to alter the assay conditions or aptamer sequence slightly to weaken the aptamer structure in the absence of ligand for use in ARTIST. Alternatively, revisiting the pool of functional sequences from the initial selection process may help with finding compatible aptamers ^52^. Further, it would be interesting to explore whether aptamers that form non-canonical structures^53^ other than G-quadruplexes are compatible with ARTIST. Additionally, adapting ARTIST to detect proteins at picomolar or femtomolar concentrations will be crucial for many diagnostic applications. This might be achieved by adding additional layers of signal amplification, either through additional transcriptional cascades or isothermal amplification of the RNA outputs^8^.

The effects of environmental conditions such as ion concentration on aptamer affinity can limit their applicability in biosensors. We demonstrate how the ARTIST system can produce a measurement of IFN-γ concentration that is mostly insensitive to that concentration of potassium ions, despite the effects of potassium on aptamer affinity^37^. Biological samples such as serum, blood, urine, and saliva, have different salt concentrations that can confound the readout of biosensors. ARTIST, through the ability to self-calibrate in response to its environment, may potentially ameliorate this issue. The incorporation of further self-calibration, background or crosstalk subtraction^54^ or even feedback control methods such as adaptation^55^ might allow ARTIST to overcome many of these limitations, or further, even to achieve robustness levels exceeding those of many traditional affinity assays.

## Materials and Methods

### Materials

O4_f, O1’_q, O2’_q, O3’_q, and O4’_q were purchased from Integrated DNA Technologies (IDT) with HPLC purification (Supplementary Table 1). All other oligonucleotides were purchased under standard desalting conditions. Triphosphates (NTPs) were purchased from ThermoFisher Scientific (EP0113). T7 RNAP was purchased in bulk (300,000 U) from Cellscript (200 U μl^−1^, C-T7300K) as well as from ThermoFisher Scientific (200 U μl^−1^, EP0113). NEB RNAPol reaction buffer (M0251S; 10X) and yeast inorganic pyrophosphatase (YIPP; M2403S; 0.1 U μl^−1^) were purchased from New England Biolabs (NEB). RNase H was purchased from ThermoFisher Scientific (EN0201; 5 U μl^−1^).

Recombinant Human IFN-γ (285-IF), IL-6 (206-IL), TNF-α (210-TA), and VEGF 165 (293-VE) were all purchased from R&D Systems Inc. (U.S.A.) in lyophilized form. Recombinant Human IFN-γ was reconstituted in sterile, deionized water, whereas Recombinant Human IL-6, TNF-α, and VEGF were all reconstituted in sterile PBS containing 0.1% BSA. Human α-thrombin (HCT0020) was purchased from Haematologic Technologies, Inc. (Essex, VT) and dissolved in 50% H_2_O/glycerol.

### dART annealing and preparation

dARTs were assembled by annealing their three strands, a promoter non-template strand (Prom-dART-nt), a non-template strand encoding the output sequence of choice (i.e. O1-dART-nt, O2-dART-nt, O3-dART-nt) and a template strand that contains the aptamer domain of choice that is complementary to the two non-template strands (i.e. IFN-O1-dART-t, Thr-O1-dART-t, IL6-O1-dART-t, TNF-O1-dART-t) at equimolar concentrations. As an example, Prom-dART-nt, O1-dART-nt, and IFN-O1-dART-t were combined in a standard 200 μL PCR tube (VMR; 20170-010) at concentrations of 1 μM per strand in 1X NEB RNAPol reaction buffer supplemented with KCl to a final concentration of 100 mM. To anneal, mixtures were heated to 90 °C, incubated for 5 min, then cooled to 20 °C at a rate of 1 °C min^−1^.

### Reporter annealing and preparation

DNA reporters were prepared by diluting the fluorophore-modified DNA strand with its partially complementary quencher-modified DNA strand at a concentration of 10 μM per strand in 1X NEB RNAPol reaction buffer. The reporter mixture was heated to 90 °C, incubated for 5 min, then cooled to 20 °C at a rate of 1 °C min^−1^.

### Amplified comparator annealing and preparation

Genelet initially in a blocked state (G1O4:B1) was prepared by mixing G1O4-nt, O4-t, and B1 together in 1X NEB RNAPol reaction buffer at equimolar concentrations. The genelet mixture was heated to 90 °C, incubated for 5 min, then cooled to 20 °C at a rate of 1 °C min^−1^.

### Reaction conditions

Reactions were all conducted at 37 °C in 1X NEB RNAPol reaction buffer supplemented with KCl to a final concentration of 100 mM and NTPs (ATP, UTP, CTP, GTP) at a final concentration of 2 mM each unless otherwise stated. We included 100 mM KCl to promote the proper folding of G-quadruplex structures within aptamer domains^26^. In addition to T7 RNA polymerase, YIPP was also included in reactions (1.35 × 10^−3^ U μl^−1^) to extend the duration of the transcription reactions. The reaction conditions mentioned above are referred to as ARTIST reaction conditions. The concentrations of each of the molecules in each experiment are given in Supplementary Tables 6-22 under Supplementary Information Section 7. The total volume of the reaction was set at 25 μL for all assays.

To perform experiments, solutions containing dART templates under ARTIST reaction condition mentioned were first added to wells of a 384-well plate. Proteins were then added at the concentrations described set at a volume of 0.5 μL and incubated for 30 min to 60 min at room temperature. In Figure 3b, we also added 0.5 μL of sterile PBS containing 0.1% BSA into the assays with 10 nM of IL6-O1-dART or TNF-O1-dART reacting with 0 nM of IL-6 or TNF-α, respectively. This was done to ensure equal salt concentrations from the PBS buffer. Similarly in Figure S10, we added 0.5 μL of sterile PBS containing 0.1% BSA into all assays that did not have 100 nM of IL-6 or TNF-α.

### Data acquisition

Fluorescence readings were then taken for 10 min to 25 mins to measure minimum fluorescence values before T7 RNAP, and YIPP were added to initiate the reactions (Supplementary Information Section 1.2, Figure S2a). At the end of the experiments, 0.5 μL of a DNA strand fully complementary to o1’_q or o4’_q was mixed into each assay at a final concentration of 2.5 μM to obtain a maximum O1 or O4 DNA reporter fluorescence intensity. Fluorescence data were then normalized using Eqn 1 as follows (Figure S2b):

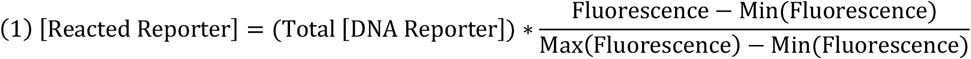

All kinetic data were obtained using either a BioTek Synergy H1 or Cytation 5 plate reader (Agilent Technologies). HEX was measured with an excitation peak of 533 nm and an emission peak of 559 nm with a gain of 80 to 100 to ensure fluorescence values were within the linear range of detection. Cy3 was measured with excitation peak of 555 nm and emission peak of 569 nm with a gain 60 to 100 to ensure fluorescence values were within the linear range of detection. FAM was measured with excitation peak of 487 nm and emission peak of 527 nm with a gain of 60. Fluorescence measurements were taken every minute during reactions. For independent replicates, all experiments were conducted using the same instrument, reagents, and experimental conditions on two or three separate days.

## Acknowledgements

The authors thank Marc Ostermeier, Elizabeth Strychalski, Simon d’Oelsnitz, Moshe Rubanov, Everett Kengmana, Colin Yancey, and Lei Zhang for insightful conversations and comments on the manuscript. H.L. was supported by the Asan Foundation Biomedical Science Scholarship. S.W.S was supported by a National Research Council Postdoctoral Fellowship. R.S. acknowledges support from NIH R21CA251027-01A1, NSF CIF Medium 2107246, and ARO award W911NF2010057. The National Institute of Standards and Technology notes that certain commercial equipment, instruments, and materials are identified in this paper to specify an experimental procedure as completely as possible. In no case does the identification of particular equipment or materials imply a recommendation or endorsement by NIST, nor does it imply that the materials, instruments, or equipment are necessarily the best available for the purpose.

## Author Contributions

H.L., S.W.S., and R.S. designed the research. H.L. conducted most of the experiments and simulations. T.X. performed some experiments in Figure 2f, 2h, and 3b of the main text and Figure S4 under Supplementary Information. X. Y. performed some experiments for Figure 3e. H.L., S.W.S., and R.S. wrote the paper with feedback from the other authors.

## Code Availability

Simulations for *K*_*d,apparent*_, reacted reporter kinetics of dARTs, reacted reporter kinetics and dose-response curves of the comparator are available at: https://github.com/hlee260/ARTIST.git

