## Supplementary Information for "Plug-and-play protein biosensors using aptamer-regulated in vitro transcription"

### Table of Contents


1. **ARTIST sequences and dART design principles**
    1.1. ARTIST species and sequence analysis
    1.2. Overview of RNA transcript design
    1.3. Method for measuring RNA transcript that reacts with DNA reporter
    1.4. Optimizing the length of the insulation domain
2. **Model to predict the binding affinity of dARTs**
    2.1. Modeling mass action rate equations
    2.2. Modeling $K_{d,apparent}$ and calculating unbound dART
    2.3. $K_{d,apparent}$ determination and validation
3. **Modularity in the general dART design**
    3.1. Modular design of dART aptamer domain for different target proteins
    3.2. Different dART outputs
    3.3. Aptamers that were unsuccessfully incorporated into dARTs
4. **Analog biosensor**
5. **Digital biosensor**
    5.1. Design of the comparator
    5.2. Simulations of the comparator
    5.3. Determining IFN-O1'-dART concentration to threshold digital ON/OFF response.
    5.4. Determining concentration of IFN-γ for digital outputs of the comparator.
    5.5. Dose-response curves of the comparator are similar with varying T7 RNAP concentrations.
    5.6. Insensitivity of the comparator to potassium ion concentration.
6. **Amplified digital biosensor**
7. **Reagents and their concentrations and experimental protocols**
8. **References**

### 1. ARTIST sequences and dART design principles

#### 1.1. ARTIST species and sequence analysis

**Supplementary Table 1.** Sequences for DNA strands involved in ARTIST. All sequences are written 5' to 3'. The aptamer domains of the dARTs are shown in bold text. Toehold sequences of DNA reporters are underlined. The sequences of G1O4-nt, O4-t, A1, and B1 are identical, respectively, to the G1S1-nt, S1-t, dA1, and dB1 sequences in Schaffter et al.[1]. dART/Reporter annealing and strand preparation protocols for sequences can be found in the main text Methods. O4_f, O1'_q, O2'_q, O3'_q, and O4'_q were purchased with HPLC purification. All other DNA strands were purchased under standard desalting conditions.

| dART non-template (nt) strands | Sequence |
|---|---|
| Prom-dART-nt | TTCTAATACGACTCACTATAGGGATG |
| O1-dART-nt | CATCCCCTACATCCACATACTAATTAAC |
| O2-dART-nt | CATCCCCTACTTTCACTTCACAACATCA |
| O3-dART-nt | CATCCCTACCATCACATTCAATAATCCT |
| **dART template (t) strands** | **Sequence** |
| IFN-O1-dART-t | GTTAATTAGTATGTGGATGTAGGGGATG**TGTTAGGGGGTTGGTTTTGGGTTGG**CATCCC TATAGTGAGTCGTATTAGAA |
| Dummy-O1-dART-t | GTTAATTAGTATGTGGATGTAGGGGATG**TTCTTCTTCTTCTTCTTCTTCTTC**CATCCC TATAGTGAGTCGTATTAGAA |
| Thr-O1-dART-t | GTTAATTAGTATGTGGATGTAGGGGATG**AGTCCGTGGTAGGGCAGGTTGGGGTGACT** CATCCCTATAGTGAGTCGTATTAGAA |
| IL6-O1-dART-t | GTTAATTAGTATGTGGATGTAGGGGATG**TGGTGGCAGGAGGACTATTTATTTGCTTTTCT** CATCCCTATAGTGAGTCGTATTAGAA |
| TNF-O1-dART-t | GTTAATTAGTATGTGGATGTAGGGGATG**TGGTGGATGGCGCAGTCGGCGACAA**CATCCC TATAGTGAGTCGTATTAGAA |
| IFN-O2-dART-t | TGATGTTGTGAAGTGAAAGTAGGGGATG**TGTTAGGGGGTTGGTTTTGGGTTGG**CATCCC TATAGTGAGTCGTATTAGAA |
| IFN-O3-dART-t | AGGATTATTGAATGTGATGGTAGGGATG**TGTTAGGGGGTTGGTTTTGGGTTGG**CATCCC TATAGTGAGTCGTATTAGAA |
| KMy-O1-dART-t | GTTAATTAGTATGTGGATGTAGGGGATG**TGGGGGTTGAGGCTAAGCCGA** CATCCCTATAGTGAGTCGTATTAGAA |

| | |
|---|---|
| VEGF-O1-dART-t | GTTAATTAGTATGTGGATGTAGGGGATG**TGTGGGGGTGGACTGGGTGGGTACC** CATCCCTATAGTGAGTCGTATTAGAA |
| **DNA reporter strands** | **Sequence** |
| O1_f | /5HEX/CTACATCCACATACTA |
| O1′_q | GTTAATTAGTATGTGGATGTAG/3IAbRQSp/ |
| O2_f | /56-FAM/CTACTTTCACTTCACAA |
| O2′_q | TGATGTTGTGAAGTGAAAGTAG/3IABkFQ/ |
| O3_f | /56-FAM/TACCATCACATTCAAT |
| O3′_q | AGGATTATTGAATGTGATGGTA/3IABkFQ/ |
| O4_f | /Cy3/GACATACAGATTAACCAGACA |
| O4′_q | GTCACTGTCTGGTTAATCTGTATGTC/3IAbRQ/ |
| **Comparator circuit strands** | **Sequence** |
| O1′-dART-nt | CATCCCGTTAATTAGTATGTGGAT |
| IFN-O1′-dART-t | ATCCACATACTAATTAACGGGATG**TGTTAGGGGGTTGGTTTTGGGTTGG**CATCCCTATAGTGAGTC GTATTAGAA |
| **Amplified comparator circuit strands** | **Sequence** |
| C1-dART-nt | CATCCCTCCTTCCATGAACGCCAAACCGTGGCGACGACCTACTC |
| Ref-C1-dART-t | GAGTAGGTCGTCGCCACGGTTTGGCGTTCATGGAAGGAGGGATG**AGTCCGTGGTAGGGCAGGTTGG GGTGACT**CATCCCTATAGTGAGTCGTATTAGAA |
| C1′-dART-nt | CATCCCAAAGAGTAGGTCGTTTCATGGAAGGA |
| IFN-C1′-dART-t | TCCTTCCATGAAACGACCTACTCTTTGGGATG**TGTTAGGGGGTTGGTTTTGGGTTGG**CATCCCTAT AGTGAGTCGTATTAGAA |
| G1O4-nt | TCCTTCCATGCACGCCAAACCGTGGCGACGTAATACGACTCACTATAGGGAGATTCGTCTCCCGAC ATACAGATTAACCAGACAGTGAC |
| A1 | TCCAGCTCTATTACGTCGCCACGGTTTGGCGTGCA |
| O4-t | GTCACTGTCTGGTTAATCTGTATGTCGGGAGACGAATCTCCCTATAGTGAGTCG |
| B1 | GAGTAGGTCGTCGCCACGGTTTGGCGTGCATGGAAGGA |
| O4-dART-nt | CATCCCGACATACAGATTAACCAGACAGTGAC |
| Ref-O4-dART-t | GTCACTGTCTGGTTAATCTGTATGTCGGGATG**AGTCCGTGGTAGGGCAGGTTGGGGTGACT**CATCC CTATAGTGAGTCGTATTAGAA |
| O4′-dART-nt | CATCCCGTCACTGTCTGGTTAATCTGTA |

| | |
|---|---|
| IFN-O4′-dART-t | TACAGATTAACCAGACAGTGACGGGATG**TGTTAGGGGGTTGGTTTTGGGTTGG**CATCCCTATAGTG AGTCGTATTAGAA |

### 1.2. Overview of RNA transcript design

The *i* and *i′* domains in the encoded transcripts of dARTs were designed so that they would hybridize, thereby creating a hairpin with a loop that sequesters the domain of the transcript whose sequence is complementary to the aptamer domain (apt') inside. Such a loop should disfavor undesired RNA secondary structure formation or other interactions between the aptamer sequence of the dART and the RNA output sequence, ensuring that the output reacts reliably downstream. We tested this design concept using the IFN-O1 dART's RNA output. Without the *i′* domain, the O1 domain of the transcript output may interact with the *i* domain and form undesirable secondary structures. With the *i′* domain, NUPACK predicts that the only the *i* and *i′* domains hybridize (Figure S1).

To predict the secondary structures of the RNA transcripts in Figure S1 and elsewhere in this work, we used NUPACK 3.2.2[2] with a temperature of 37 °C and the default salt conditions (1 M $Na^+$, 0 M $Mg^{2+}$). Although there is 6 mM $MgCl_2$ in our transcription buffer, there is a total of 8 mM of NTPs (2 mM of each ATP, UTP, CTP, GTP), which will sequester $Mg^{2+}$, so the concentration of free $Mg^{2+}$ is unknown. NUPACK cannot be used to predict G-quadruplex formation (as this is tertiary rather than secondary structure), so we assume that all single-stranded aptamer domains fold into a G-quadruplex structure due to 100 mM of K+ under ARTIST reaction conditions[3,4].

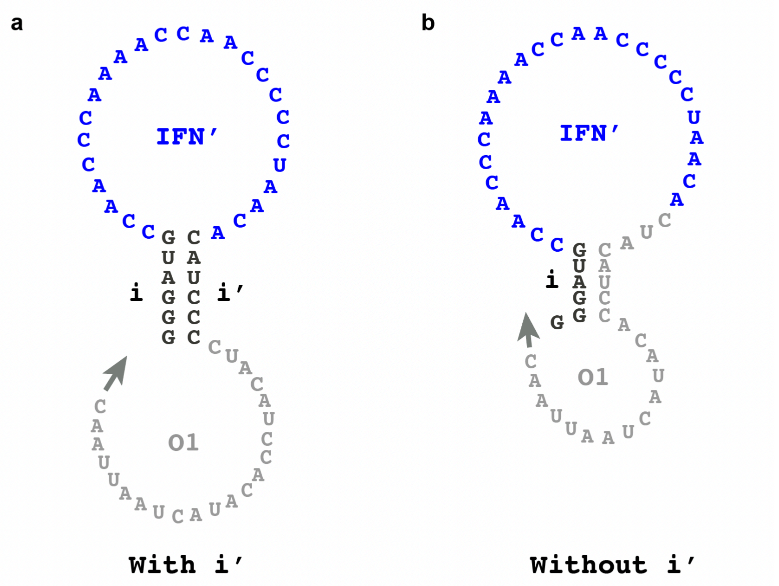


**Figure S1.** NUPACK[2] predicts that insulator domains prevent the formation of unintended secondary structure within dART RNA outputs. **(a)** A transcript from IFN-O1-dART[5] containing a *i'* domain is predicted to form a hairpin structure, leaving the output (O1) domain free. **(b)** undesired secondary structure forms in a transcript from a dART without the *i'* domain: bases from the O1 domain hybridize with *i*. This secondary structure could prevent or alter the rate of the output's participation in downstream reactions such as strand displacement.

#### 1.3. Method for measuring the amount of RNA transcript that reacts with DNA reporter

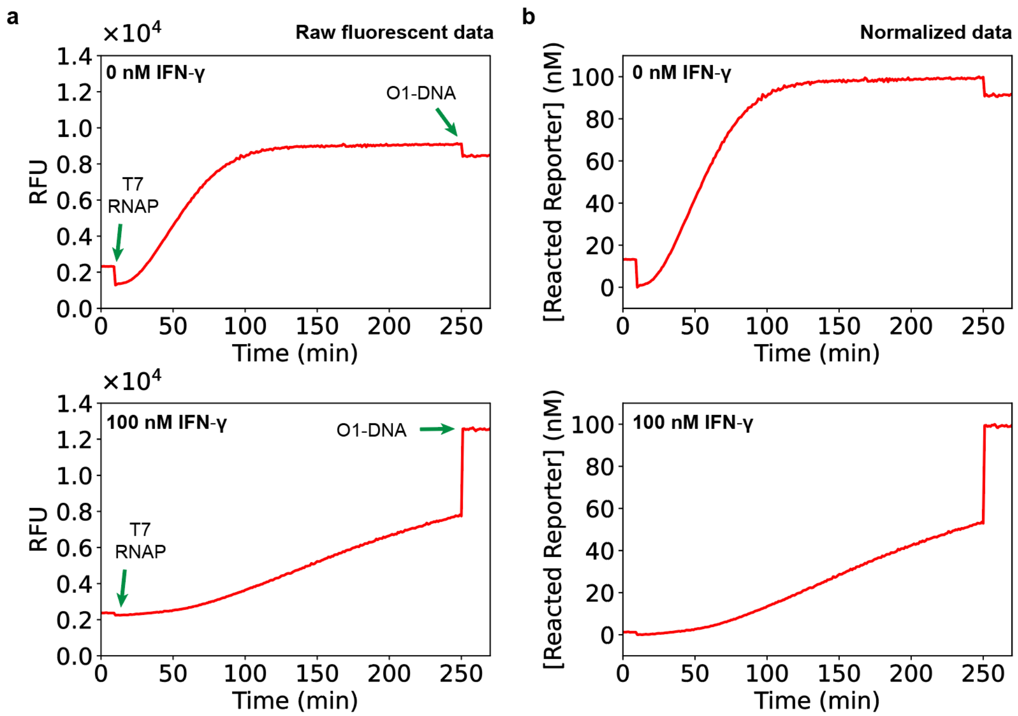


**Figure S2.** Normalization of reacted reporter measurements. **(a)** Representative examples of how time-dependent measurements of reacted reporter concentration were obtained from fluorescence measurements. 10 nM of IFN-O1-dART, 100 nM of O1 DNA Reporter, 0 nM or 100 nM of IFN-γ are initially combined (see main text Methods). Fluorescence readings were first taken for 10 min. 2 U $\mu L^{-1}$ of T7 RNAP was then added. After measuring fluorescence for another 240 min to 360 min, 2.5 μM of O1-DNA, which is fully complementary to o1'_q, was mixed into each sample to obtain an internal maximum DNA reporter fluorescence value. We waited at least 10 min after adding in O1-DNA to measure this maximum fluorescence. Since the reacted fluorescent signal is proportional to the extent of the reaction[8], the concentration of reacted reporter could be obtained from the fluorescence using the Eqn 1 in the main text Methods. **(b)** The concentrations of reacted reporter over time for reactions containing 10 nM of IFN-O1-dART and 100 nM of O1 DNA Reporter with either 0 nM (top) or 100 nM (bottom) of IFN-γ.

#### 1.4. Optimizing the length of the insulation domain

For transcription factor proteins that repress transcription, the number of bases between the T7 promoter and the beginning of the repressor's operator sequence on the DNA template greatly influences transcription and repression efficiency[6–9]. We hypothesized that the number of bases between the T7 promoter and the beginning of the aptamer domain on a dART could also influence aptamer-regulated transcription efficiency and the difference in unbound and protein-bound dART transcription rates. To explore this possibility, we designed a series of dARTs in which the length of the dsDNA domain between the promoter and aptamer domains was varied between 2 bases and 22 bases. All these dART variants were still designed to produce a transcript with a 6 base hairpin to insulate the sequence transcribed from the aptamer domain from the output sequence[1]. This required introducing an *h* domain that was complementary to a portion of the apt' sequence in the transcripts of the dART variants having fewer than 6 bases between the promoter and the aptamer domain (Figure S3 and Supplementary Table 2). For dARTs with more than 6 bases between the promoter and the aptamer domain, only the length of the sequence upstream of the insulation domain was extended, termed the leader, or *l*, domain (Figure S4 and Supplementary Table 2).

We characterized the reacted reporter kinetics for the dART variants both in the presence and absence of IFN-γ. dART variants with greater than 6 bases between the promoter and aptamer domain had slow reacted reporter kinetics even in the absence of IFN- γ. dART variants with 2 bases, 4 bases, or 6 bases between the promoter and aptamer domain had fast reacted reporter kinetics in the absence of IFN-γ, and only the 6 base variant had a substantial repression of reacted reporter signal with IFN-γ (Figure S5). Based on these results, all the dARTs in this study were designed with 6 bases between the promoter and aptamer domains.

**Supplementary Table 2:** DNA strands and sequences involved in optimizing the length between the promoter and the aptamer domains of dARTs. The aptamer domain is shown in bold. The $i_2$, $i_4$, $i_6$ domains are underlined and designate lengths of 2 bases, 4 bases, and 6 bases, respectively The sequences appended to $i_6$ (*+4, +8, +12, +16*) are the *l* domains, which are shown in orange. The *h* domain, which is in teal font, is partially complementary to the aptamer domain so that the encoded transcript of dARTs with (2 to 4) base insulation domain lengths form a hairpin that prevents the O1 domain from forming undesired secondary structures. The portions of the aptamer sequence complementary to the *h* domain are both bolded as well as highlighted in cyan. dART variants with greater than 6 bases between the promoter and aptamer were prepared using O1-dART-nt-i_6 for the output nt (non-template) strands.

| Promoter nt | Sequence |
|---|---|
| Prom-dART-nt-i2 | TTCTAATACGACTCACTATA GG |
| Prom-dART-nt-i4 | TTCTAATACGACTCACTATA GGGA |
| Prom-dART-nt-i6 | TTCTAATACGACTCACTATA GGGATG |
| Prom-dART-nt-i6+4 | TTCTAATACGACTCACTATA GGGA GGGATG |

| | |
|---|---|
| Prom-dART-nt-i6+8 | TTCTAATACGACTCACTATA GGGAGATT GGGATG |
| Prom-dART-nt-i6+12 | TTCTAATACGACTCACTATA GGGAGATTCGTC GGGATG |
| Prom-dART-nt-i6+16 | TTCTAATACGACTCACTATA GGGAGATTCGTCTCCC GGGATG |
| **Output nt** | **Sequence** |
| O1-dART-nt-i_2 | TTGG CC TACATCCACATACTAATTAAC |
| O1-dART-nt-i_4 | GG TCCC TACATCCACATACTAATTAAC |
| O1-dART-nt-i_6 | CATCCC TACATCCACATACTAATTAAC |
| **dART template** | **Sequence** |
| IFN-O1-dART-t-i2 | GTTAATTAGTATGTGGATGTA GG CCAA **TGTTAGGGGGTTGGTTTTGGGTTGG** CC TATAGTGAGTCGTATTAGAA |
| IFN-O1-dART-t-i4 | GTTAATTAGTATGTGGATGTA GGGA CC **TGTTAGGGGGTTGGTTTTGGGTTGG** TCCC TATAGTGAGTCGTATTAGAA |
| IFN-O1-dART-t-i6 | GTTAATTAGTATGTGGATGTA GGGATG **TGTTAGGGGGTTGGTTTTGGGTTGG** CATCCC TATAGTGAGTCGTATTAGAA |
| IFN-O1-dART-t-i6+4 | GTTAATTAGTATGTGGATGTA GGGATG **TGTTAGGGGGTTGGTTTTGGGTTGG** CATCCC TCCC TATAGTGAGTCGTATTAGAA |
| IFN-O1-dART-t-i6+8 | GTTAATTAGTATGTGGATGTA GGGATG **TGTTAGGGGGTTGGTTTTGGGTTGG** CATCCC AATCTCCC TATAGTGAGTCGTATTAGAA |
| IFN-O1-dART-t-i6+12 | GTTAATTAGTATGTGGATGTA GGGATG **TGTTAGGGGGTTGGTTTTGGGTTGG** CATCCC GACGAATCTCCC TATAGTGAGTCGTATTAGAA |
| IFN-O1-dART-t-i6+16 | GTTAATTAGTATGTGGATGTA GGGATG **TGTTAGGGGGTTGGTTTTGGGTTGG** CATCCC GGGAGACGAATCTCCC TATAGTGAGTCGTATTAGAA |

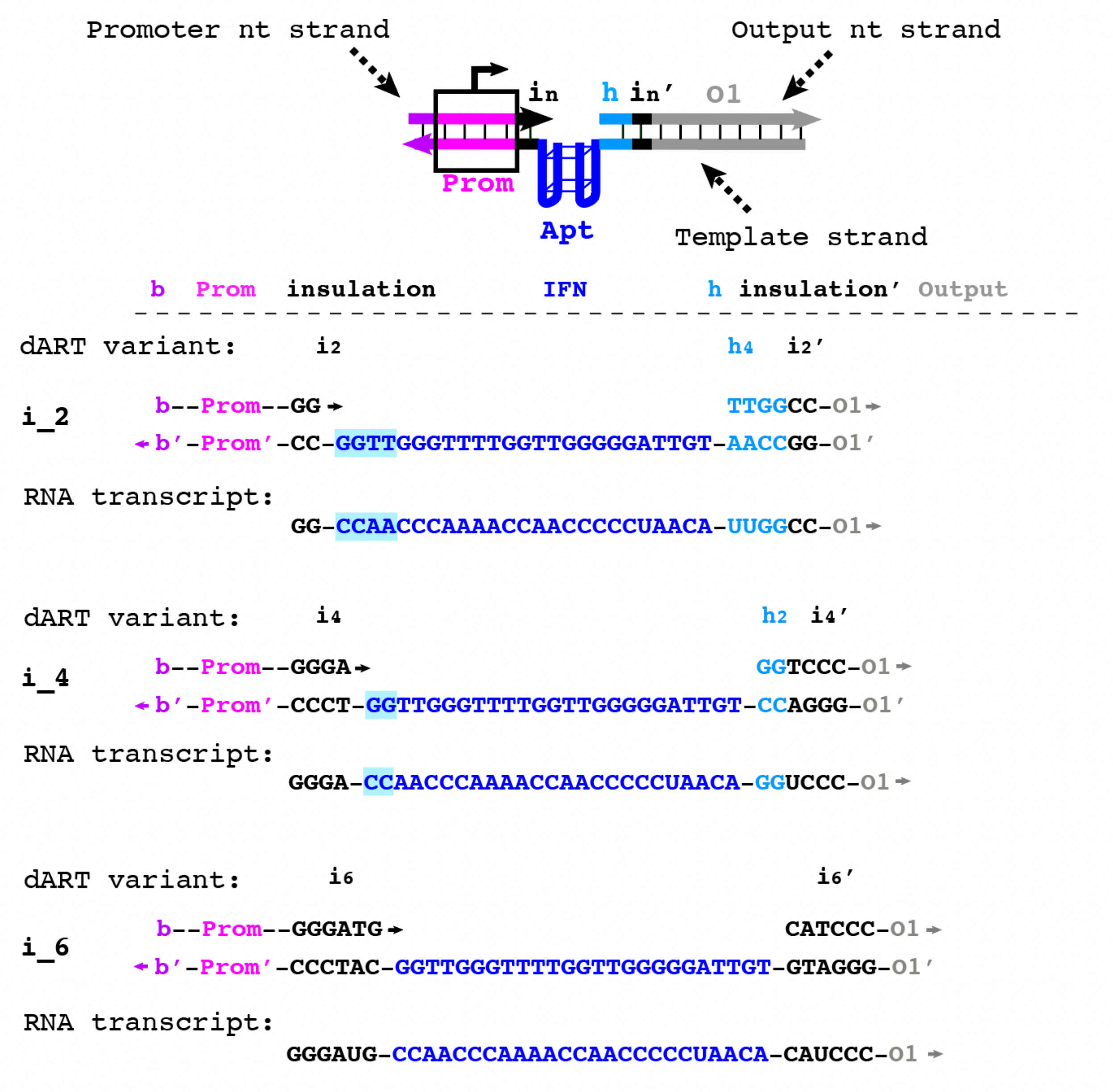


**Figure S3.** Schematic of sequences and domains for the dART variants and their RNA transcripts with $i_2$, $i_4$, and $i_6$ between the promoter and aptamer domains. Sequences highlighted in teal can hybridize to the *h* domain to form a hairpin. The *h* domain is in teal font. The i_6 dART variant was used to design all other dARTs in this study.

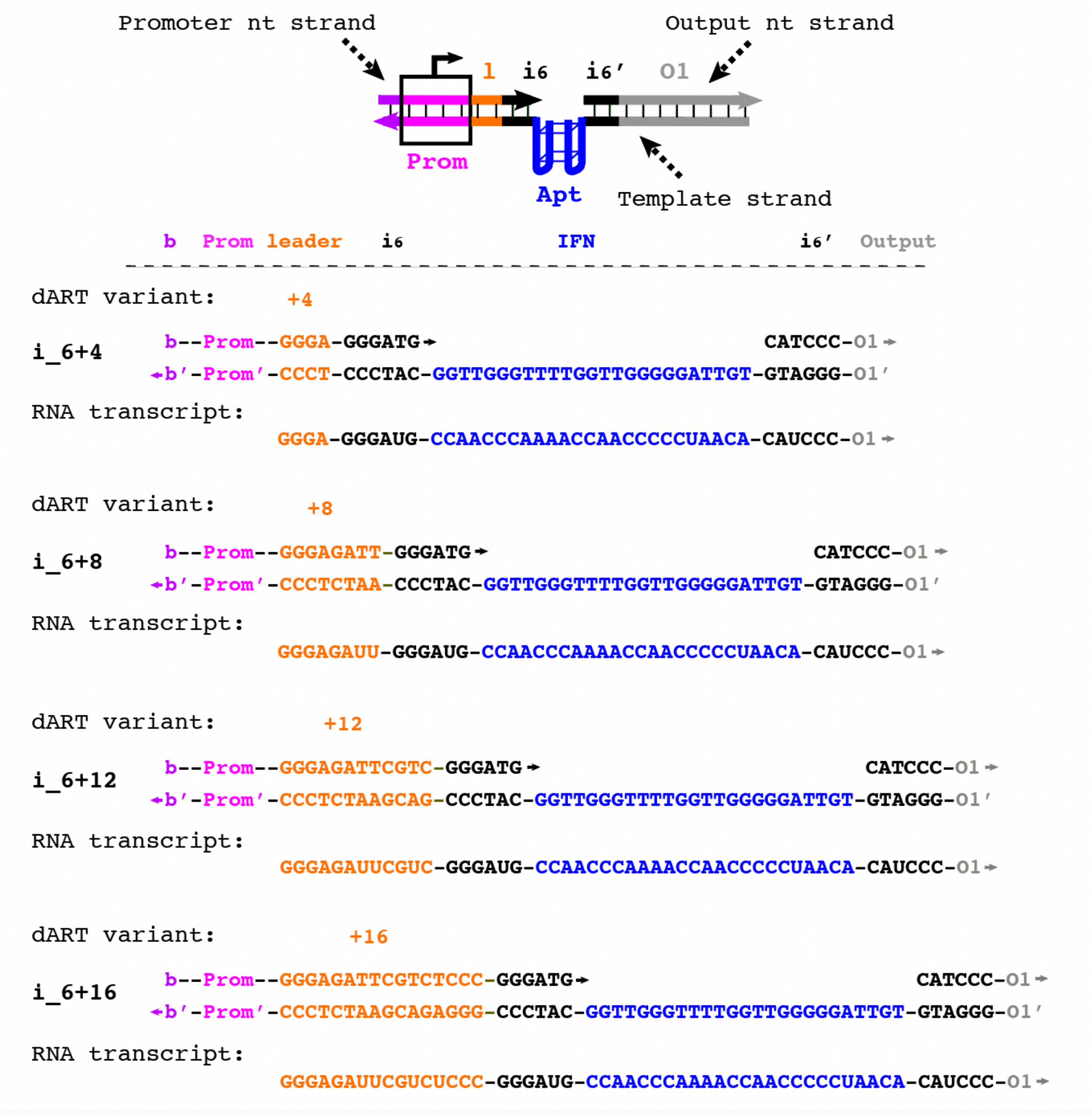


**Figure S4:** Schematic of sequences and domains for the dART variants and their RNA transcripts with *leader* domains of varying length (*+4, +8, +12,* and *+16*) and $i_6$ between the promoter and aptamer domains. *Leader* domains are in orange font.

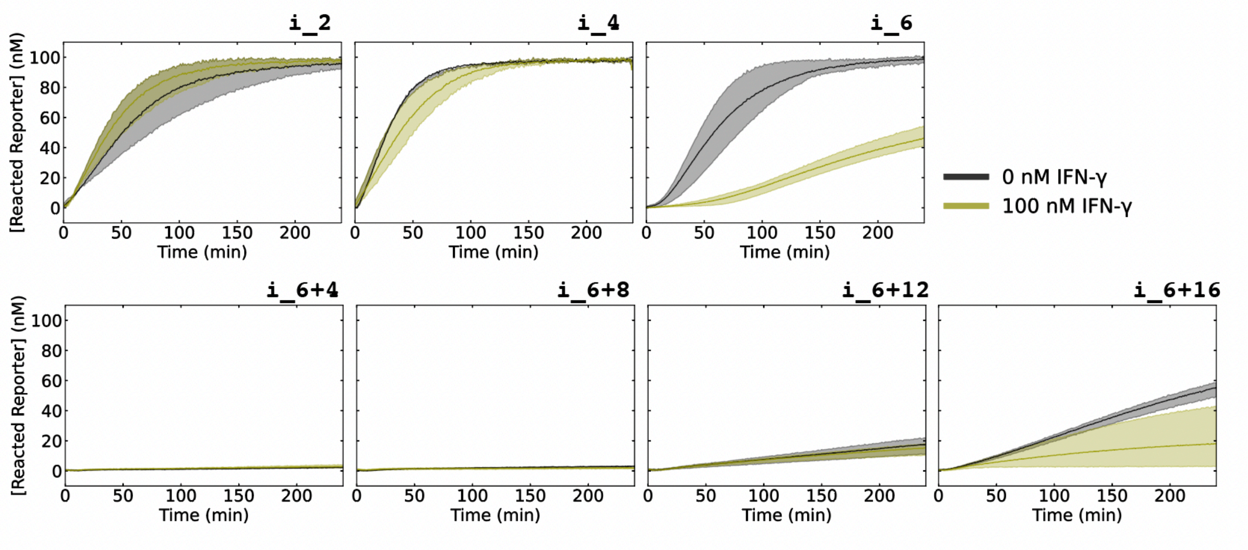


**Figure S5.** Reacted reporter kinetics induced by dARTs with varying lengths between the promoter and aptamer (2 to 22) bases and 0 nM or 100 nM IFN-γ. For all measurements, we used 10 nM dART and 100 nM reporter. Shaded regions denote minimum/maximum values of reacted reporter concentrations for three independent replicates.

### 2. Kinetic modeling of dARTs

#### 2.1. Modeling mass action rate equations

We used a mass action kinetic model to predict changes in reacted reporter concentration over time. In the model, RNA is transcribed from an unbound dART and can bind to unreacted reporter (Figure S6):

$$(S1)\ \frac{d[RNA]}{dt} = k_{txn}[dART]_{unbound} - k_{sd}[RNA][Reporter]_{unreacted},$$

$$(S2)\ \frac{d[Reporter]_{reacted}}{dt} = k_{sd}[RNA][Reporter]_{unreacted}.$$

All simulations of Eqns S1 and S2 were conducted using Python's SciPy package (version 1.10.1) using the odeint library. In Eqns S1 and S2, $[RNA]$ is the concentration of free RNA transcript, $[dART]_{unbound}$ is the concentration of the unbound dART, $[Reporter]_{unreacted}$ is the concentration of the unreacted DNA reporter, and $[Reporter]_{reacted}$ is the concentration of reacted reporter. The rate of RNA production in the model is $k_{txn}[dART]_{unbound}$. $k_{txn}$ is an apparent first order rate constant of transcription. $[dART]_{unbound}$ is determined using Eqn S3 (see Section 2.2). The strand displacement rate constant ( $k_{sd}$ ) was assumed to be $1 \times 10^5\ M^{-1}s^{-1}$, which is consistent with previous measurements[10] of rate constants for strand displacement with a toehold length of 6 bases. $k_{txn}$ was fit as $0.0015\ s^{-1}$ using the experimentally measured reacted reporter kinetics when 10 nM IFN-O1-dART and 0 nM IFN-γ were present (Figure S7).

In our experiments, we incubate the dARTs with their ligands for 30-60 min prior to adding T7 RNAP. In the model, we assumed that dARTs and their ligands reach an equilibrium between bound and unbound dARTs that are determined by the apparent dissociation constant ($K_{d,apparent}$) between the protein and the aptamer and the protein concentration[11] (as discussed below in Supplementary Section S2.2 and Equation S3) during this initial incubation time.

We also assume that when a ligand is bound to the dART's aptamer's domain, the dART cannot be transcribed.

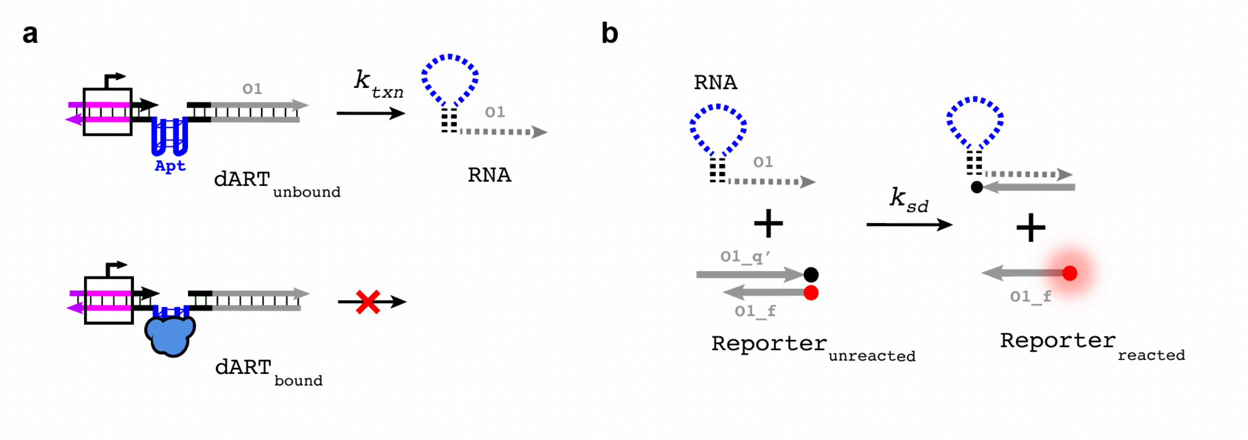


**Figure S6.** Schematic of reactions in our kinetic model. **(a)** An unbound dART can be recognized by T7 RNAP which transcribes an output with a rate constant of $k_{txn}$. However, a dART bound to its protein ligand cannot undergo transcription to produce RNA. **(b)** The RNA transcript produced from an unbound dART can bind to the 6-base toehold on a DNA Reporter complex and undergo a toehold-mediated strand displacement reaction with a rate constant of $k_{sd}$. The measured output of this reaction is $[Reporter]_{reacted}$, the concentration of a fluorescent DNA strand.

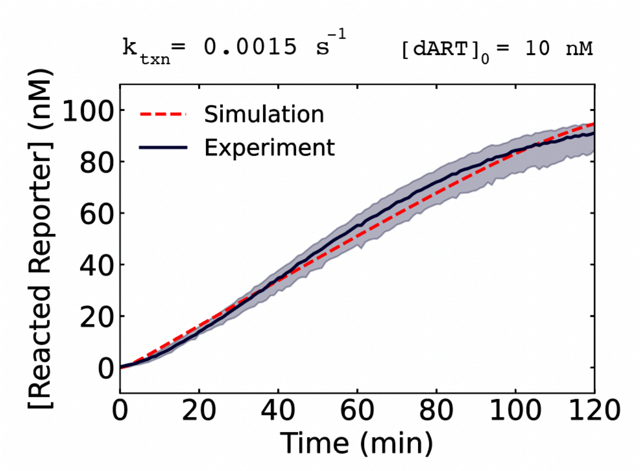


**Figure S7.** Fitting the transcription rate of 10 nM IFN-O1-dART and 0 nM of IFN-γ. A $k_{txn}$ value of 0.0015 $s^{-1}$ was fit to the average transcription plot that was measured experimentally. Shaded regions represent minimum/maximum values of reacted reporter concentration for three independent replicates.

### 2.2. Modeling $K_{d,apparent}$ and calculating unbound dART

We assume that unbound dARTs can bind to their protein ligand with a binding affinity defined by its dissociation constant ($K_d$):

$$(S3)\ K_d = \frac{[L][dART]_{unbound}}{[L{:}dART]},$$

where $[L]$ and $[L{:}dART]$ are the concentrations of free ligand and ligand-bound dART, respectively.

Based on the assumption that the binding reaction between the aptamer and the dART is at equilibrium, we can use Eqn S3 to compute $[dART]_{unbound}$ for a given initial dART concentration ($[dART]_0$) and initial ligand concentration ($[L]_0$) in terms of the dissociation constant. From mass balances[12] $[L]_0 = [L] + [L{:}dART]$ and $[dART]_0 = [dART]_{unbound} + [L{:}dART]$, we can express Eqn S3 in terms of the initial concentrations $L$ and dART and the bound dART concentration:

$$(S4)\ K_d = \frac{([L]_0 - [L{:}dART])(\,[dART]_0 - [L{:}dART])}{[L{:}dART]}.$$

We can rearrange Eqn S4 and solve for $[L{:}dART]$ using the quadratic formula:

$$(S5)\ [L{:}dART] = \frac{[dART]_0 + [L]_0 + K_d - \sqrt{([L]_0 + [dART]_0 + K_d)^2 - 4[dART]_0[L]_0}}{2}$$

For the $K_{d,apparent}$ value for a given dART, we can use Eqn S5 and $[dART]_{unbound} = [dART]_0 - [L{:}\,dART]$ to predict the equilibrium concentrations of unbound IFN-O1-dART from $[dART]_0$ (in our experiments $[dART]_0$ was 10 nM or 1 nM) for different initial concentrations of IFN-γ, *i.e.,* $[L]_0$ (in our experiments, this concentration was varied between 0 nM to 1000 nM).

#### 2.3. $K_{d,apparent}$ determination and validation

Using Eqns S3, S4, and S5, we fit $K_{d,\mathrm{apparent}}$, to the measured $[Reporter]_{reacted}$ values after 120 min for 10 nM of IFN-O1-dART in the presence of (0, 5, 10, 25, 50, 100, 250, 500, or 1000) nM of IFN-γ (Figure 2e of the main text, Supplementary Table 3). The value of $K_{d,apparent}$ fit was 8 nM.

Kinetic simulations using the fit $K_{d,apparent}$ exhibited good agreement with experimental measurements (Figure 2f of the main text). For comparison, $K_{d,apparent}$ values 5-fold higher or lower than the fit value of 8 nM did not produce a close correspondence with experimental results (Figure S8). The $K_{d,apparent}$ value of 8 nM was also consistent with experiments using a lower initial concentration of dART (Figure 2h and 2i in main text).

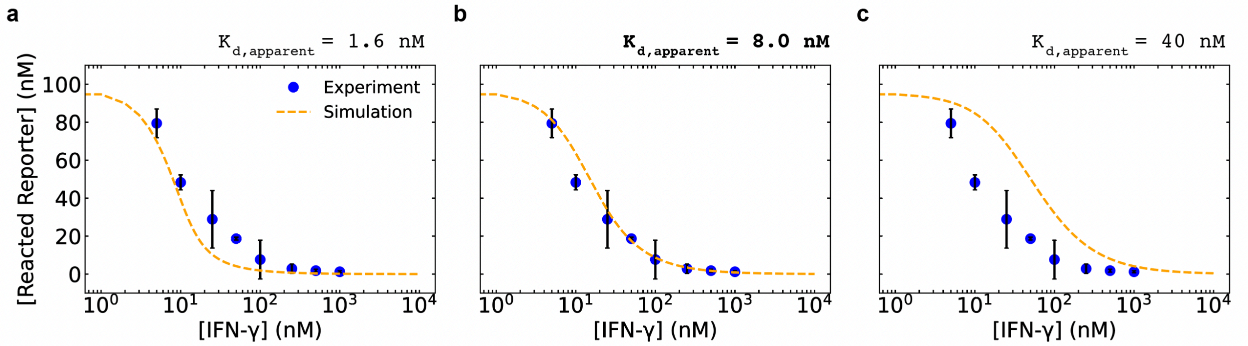


**Figure S8.** Simulated and experimental dose-response curves for 10 nM of IFN-O1-dART detecting 0 nM to 100 nM of IFN-γ with $K_{d,apparent}$ of **(a)** 1.6 nM, **(b)** 8 nM, and **(c)** 40 nM. Based on our results, 8 nM is the most suitable value of $K_{d,apparent}$ between IFN-O1-dART and IFN-γ to fit the observed dose-response behavior. For **(a)**-**(c)**, error bars show the standard deviations of three independent replicates. The simulated and experimental dose-response curves for $K_{d,apparent}$ of 8 nM are also shown in the main text Figure 2e.

**Supplementary Table 3.** Predicted unbound IFN-O1-dART concentrations that correspond to 0 nM to 1000 nM IFN-γ and rate constants for the simulations presented in Figure 2f and 2g of the main text. The unbound IFN-O1-dART concentration are calculated as described in Section 2.2 using a $K_{d,apparent}$ value of 8 nM, an initial dART concentration of 10 nM, and initial IFN-γ concentrations of (0, 5, 10, 25, 50, 100, 250, 500, or 1000) nM.

| [IFN-γ] (nM) | [IFN-O1-dART] (nM) | Rate Parameters | Rate constants |
|---|---|---|---|
| 0 | 10.0 | k_txn | 0.0015 $s^{-1}$ |
| 5 | 7.6 | k_sd | $1\times10^{5}$ $M^{-1}s^{-1}$ |
| 10 | 5.8 | | |
| 25 | 3.1 | | |
| 50 | 1.6 | | |
| 100 | 0.8 | | |
| 250 | 0.3 | | |
| 500 | 0.2 | | |
| 1000 | 0.1 | | |

**Supplementary Table 4.** Predicted unbound IFN-O1-dART concentrations that correspond to 0 nM to 100 nM IFN-γ and rate constants for the simulations presented in Figure 2h and 2i of the main text. Unbound IFN-O1-dART concentrations were calculated as described in Section 2.2 using a $K_{d,apparent}$ value of 8 nM, initial dART concentration of 10 nM, and initial IFN-γ concentrations of (0, 1, 3, 5, 10, or 100) nM.

| [IFN-γ] (nM) | [IFN-O1-dART] (nM) | Rate Parameters | Rate constants |
|---|---|---|---|
| 0 | 1.0 | k_txn | 0.0046 $s^{-1}$ |
| 1 | 0.9 | k_sd | $1\times10^{5}$ $M^{-1}$ $s^{-1}$ |
| 3 | 0.7 | | |
| 5 | 0.6 | | |
| 10 | 0.5 | | |
| 100 | 0.1 | | |

### 3. dART Modularity

#### 3.1. Modular design of dART aptamer domains for different target proteins

dART schematics with modular aptamer domains and their predicted RNA secondary structures are shown in Figure S9 and S10.

To verify the selectivity of the dARTs for their corresponding proteins, we incubated 10 nM of IFN-O1-dART, Thr-O1-dART, IL6-O1-dART, and TNF-O1-dART with 100 nM of either BSA, thrombin, TNF-α, IL-6, or IFN-γ in different samples, and measured the concentration of reacted reporter after 120 min of transcription for each sample (Figure S11). Dummy-O1-dART was also combined with these sample proteins as a control. The endpoint reacted reporter concentrations in Figure S11 were used to generate the heat map in Figure 3c in the main text.

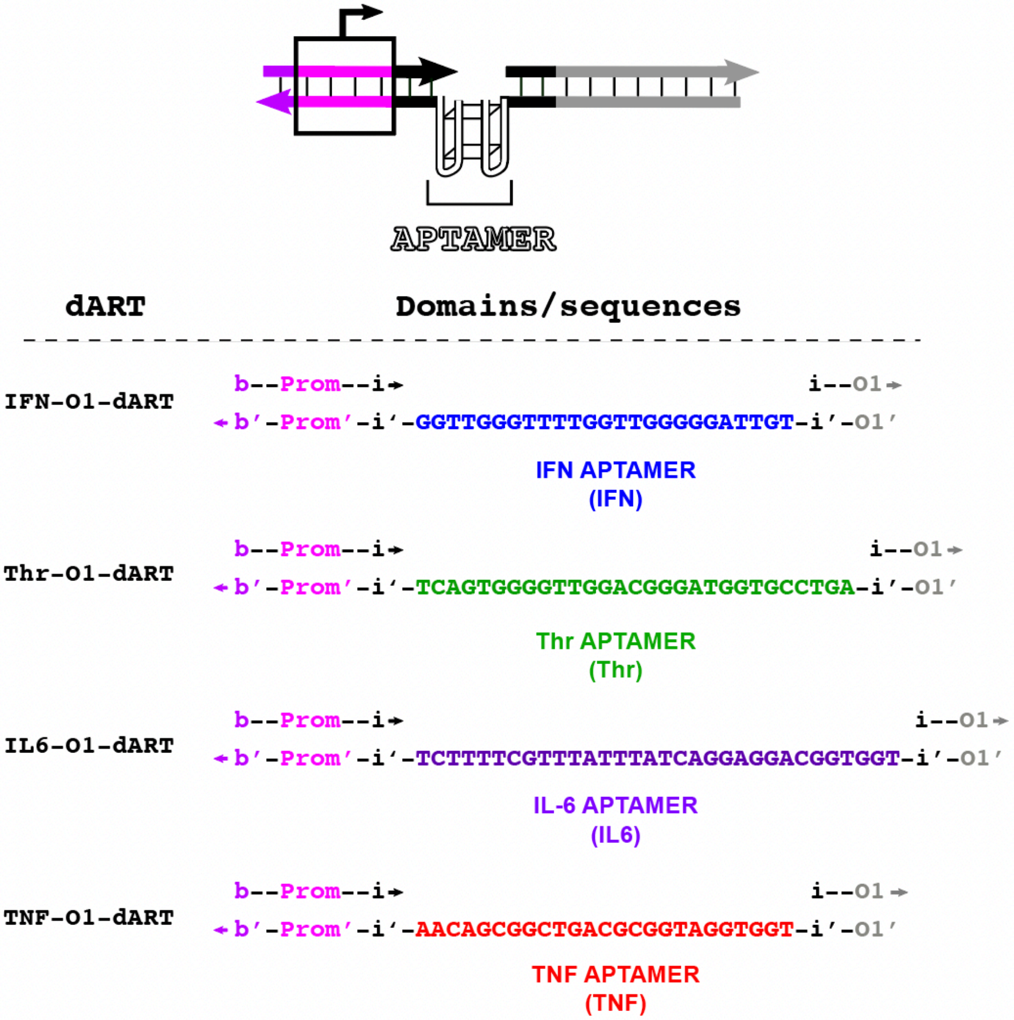


**Figure S9.** Sequences of the aptamer domains of IFN-O1-dART[5], Thr-O1-dART[13], IL6-O1-dART[14], and TNF-O1-dART[15]. Full sequences of the dART strands are provided in Supplementary Table 1.

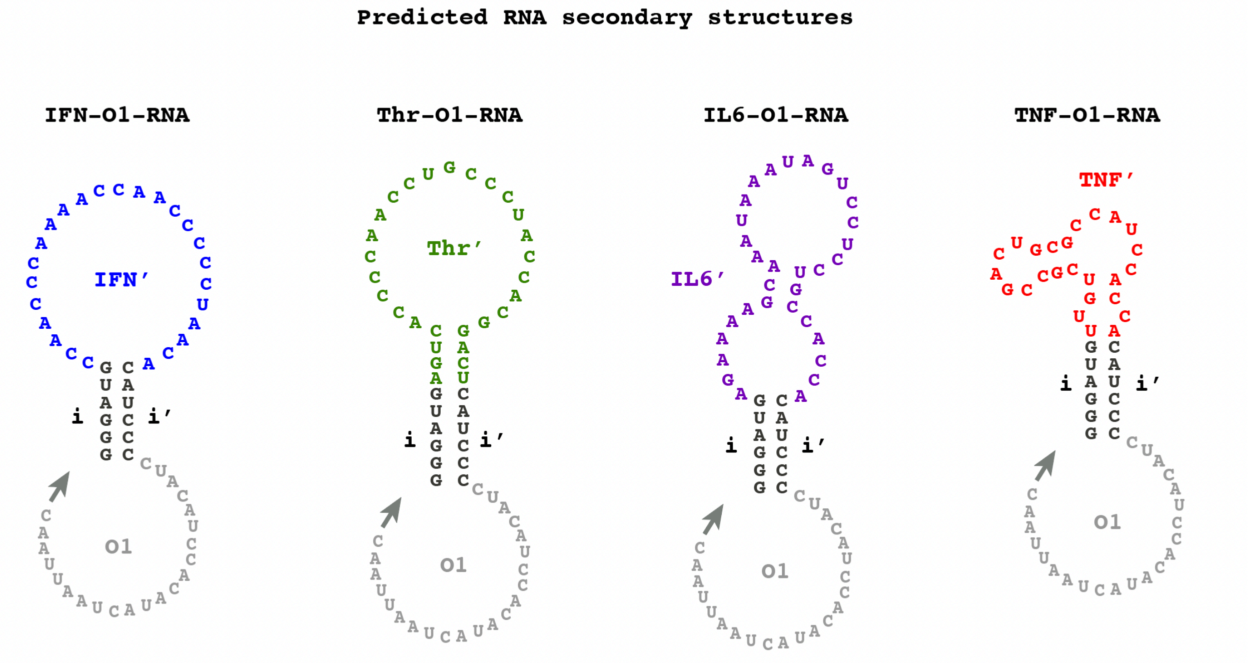


**Figure S10.** RNA secondary structures of the encoded RNA transcripts of IFN-O1-dART, Thr-O1-dART, IL6-O1-dART, and TNF-O1-dART predicted by NUPACK[2].

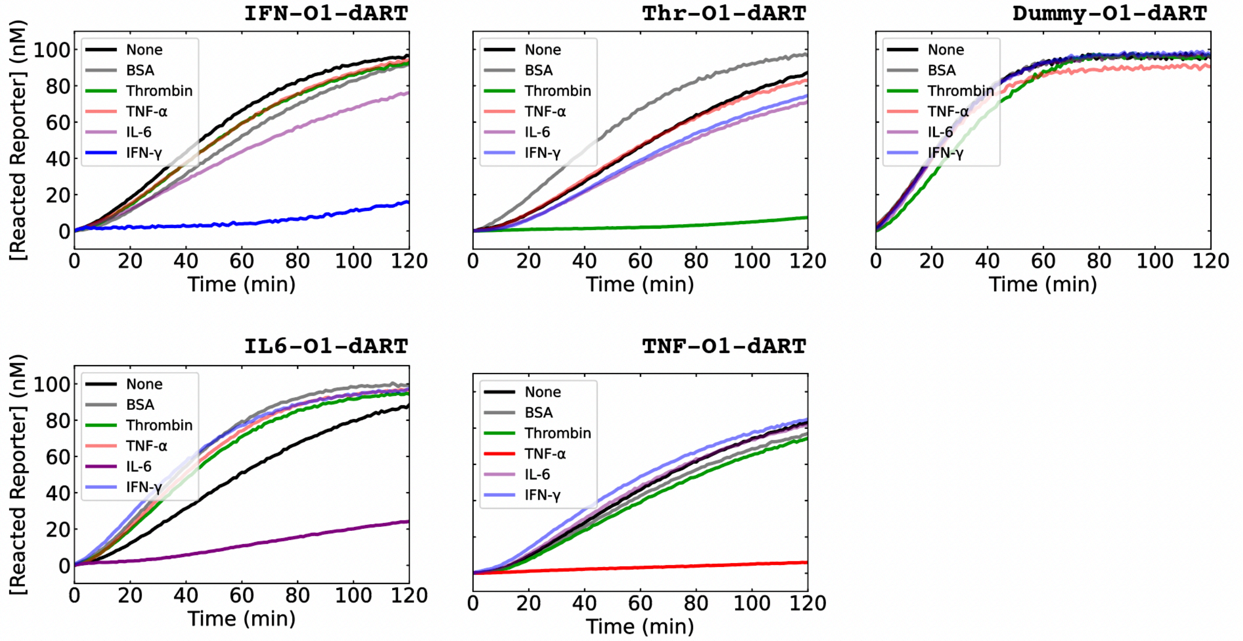


**Figure S11.** Reacted reporter kinetics for 10 nM of IFN-O1-dART, Thr-O1-dART, IL6-O1-dART, TNF-O1-dART, and a Dummy-O1-dART (a control) subjected to 100 nM of BSA, IFN-γ, thrombin, IL-6, or TNF-α. The heat map in main text Fig. 3c shows the concentration of reacted reporter after 120 min for IFN-O1-dART, Thr-O1-dART, IL6-O1-dART, TNF-O1-dART, and Dummy-O1-dART with each protein.

#### 3.2. Different dART outputs

dART schematics with modular output domains (O1, O2, O3) and their predicted RNA secondary structures are shown in Figures S12 and S13. Full sequences of the dART strands are provided in Supplementary Table 1.

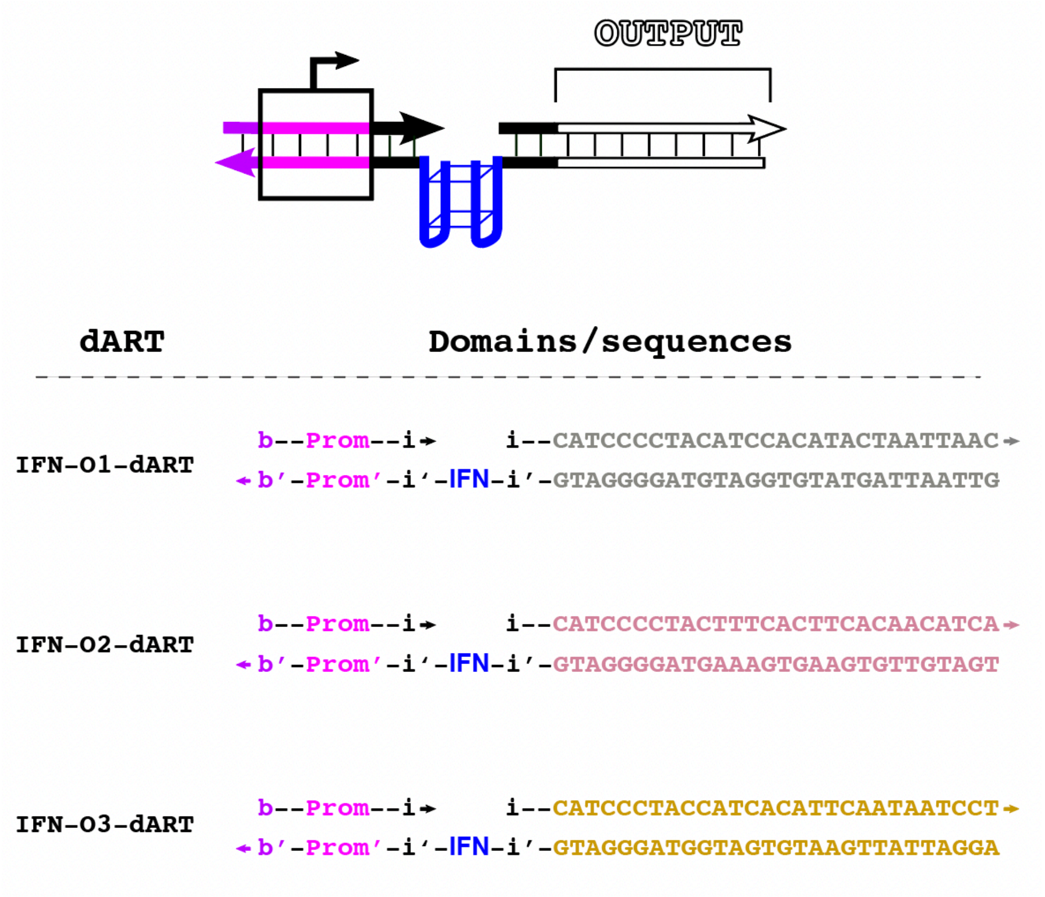


**Figure S12.** The output sequences of IFN-O1-dART, IFN-O2-dART, and IFN-O3-dART.

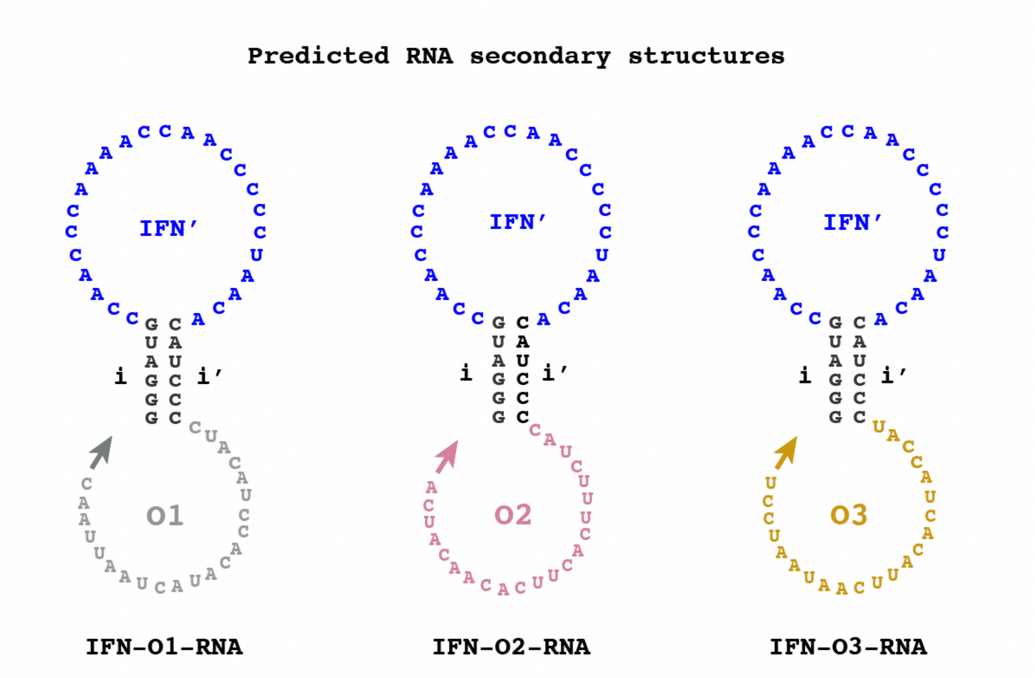


**Figure S13.** RNA secondary structures of the IFN-RNA encoded by IFN-O1-dART, IFN-O2-dART, and IFN-O3-dART predicted by NUPACK[2].

#### 3.3. Aptamers that were unsuccessfully incorporated into dARTs

We also designed a KMy-O1-dART and a VEGF-O1-dART, which had aptamers for kanamycin[16] and VEGF[17], respectively (Figure S14). We tried to embed these two aptamers into our general dART design, and used NUPACK[2] to predict the secondary structures of the RNA transcripts (Figure S15).

We first mixed 10 nM of KMy-O1-dART and VEGF-O1-dART with 100 nM O1 DNA Reporter in the presence of either 0 mM or 100 mM potassium ions to see whether the dARTs would still transcribe when the formation of G-quadruplexes was favored (Figure S16). KMy-O1-dART had fast reacted reporter kinetics in the absence of potassium, but with 100 mM KCl, the reacted reporter kinetics were minimal. This may be because the G-quadruplex structure of the kanamycin aptamer is stable enough to repress transcription even without a bound ligand. VEGF-O1-dART also showed relatively little reporter reaction regardless of 0 mM or 100 mM potassium. This could be due to unintended reactions of the RNA transcript, because it is predicted to adopt a different structure than designed (Figure S15).

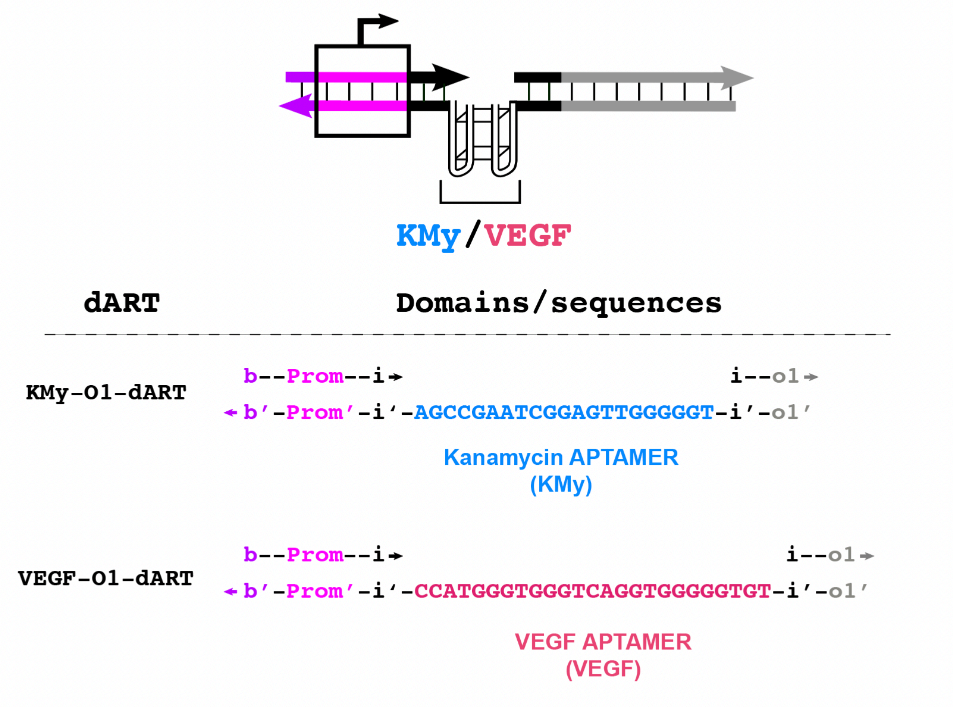


**Figure S14.** Sequences of the aptamer domains of KMy-O1-dART and VEGF-O1-dART. Full sequences of the dART strands are provided in Supplementary Table 1.

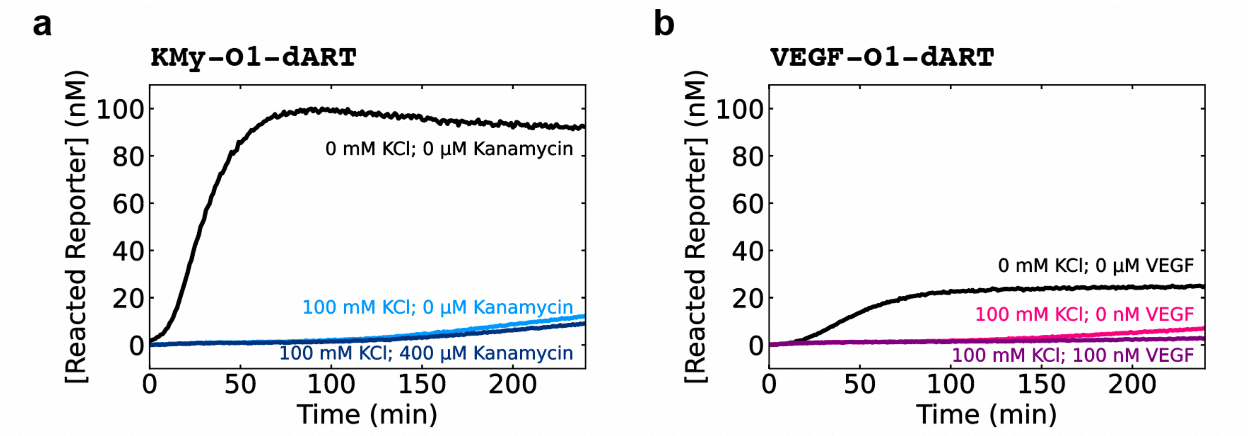


**Figure S15.** Reacted reporter kinetics for (**a**) the KMy-O1-dART and (**b**) the VEGF-O1-dART. 10 nM of KMy-O1-dART or VEGF-O1-dART was mixed with 100 nM of O1 DNA Reporter with or without KCl and with or without ligand as indicated in the plots.

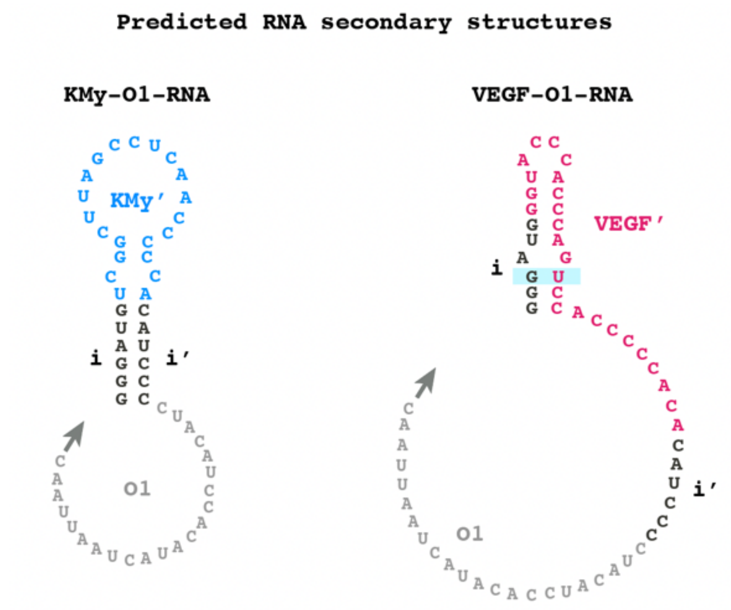


**Figure S16.** Predicted RNA secondary structures of KMy-O1-RNA and VEGF-O1-RNA. The G-U wobble in the VEGF-O1-RNA is highlighted in cyan. Secondary structure predictions of the transcripts were done using NUPACK[2].

### 4. Analog biosensor

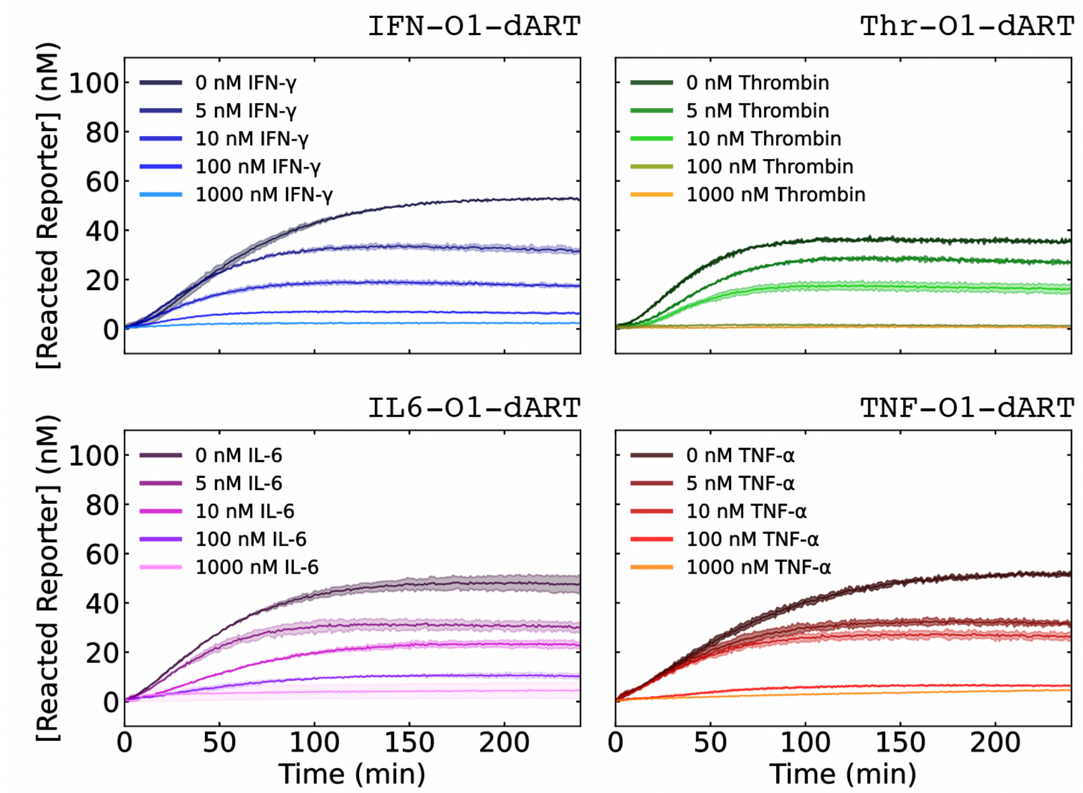


**Figure S17.** Analog biosensors in which the steady-state concentration of reacted reporter indicates protein concentration. Reacted reporter concentration over time of analog biosensors for IFN-γ, thrombin, TNF-α, and IL-6. 10 nM IFN-O1-dART, Thr-O1-dART, IL6-O1-dART, and TNF-O1-dART were mixed with 100 nM of reporter, and 0 nM to 1000 nM of their corresponding input proteins. 2 U μL$^{-1}$ of T7 RNAP and $2 \times 10^{-3}$ U μL$^{-1}$ of RNase H were used in each assay. Shaded regions enclose the minimum and maximum values of reacted reporter concentration for two replicates. The IFN-O1-dART data is also shown in main text Figure 4b. Endpoint measurements of reacted reporter concentrations at 240 min are shown in main text Figure 4c.

### 5. Digital biosensor

#### 5.1. Design of the comparator

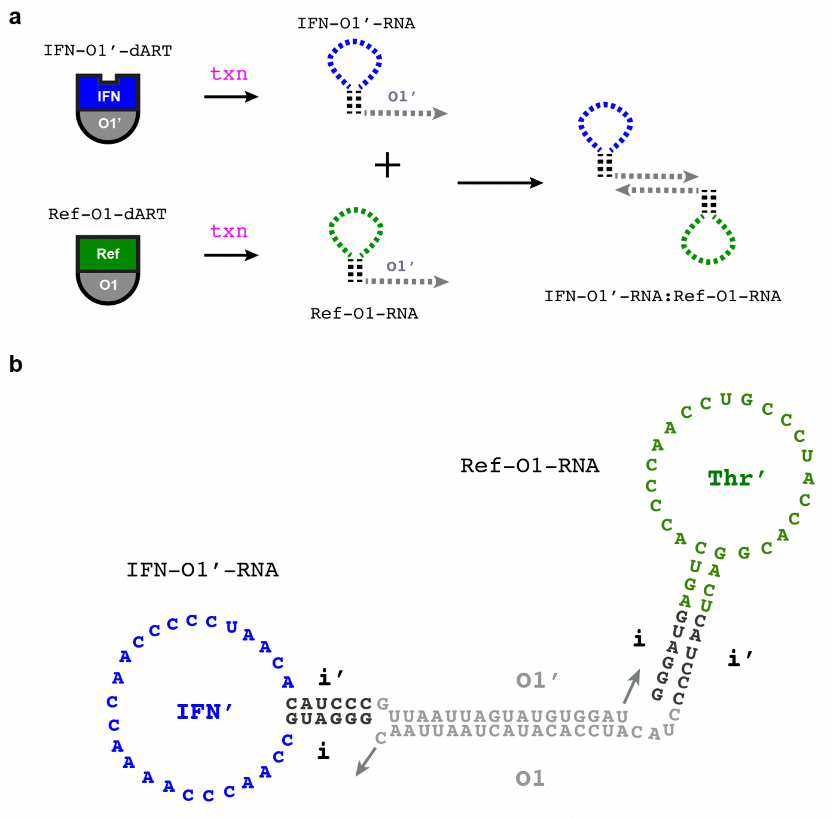


**Figure S18.** Reaction between IFN-O1'-RNA and Ref-O1'RNA. **(a)** When IFN-γ is not bound to IFN-O1'-dART, IFN-O1'-RNA is produced and can hybridize with Ref-O1-RNA. **(b)** Secondary structure of the IFN-O1'-RNA / Ref-O1-RNA complex predicted by NUPACK. The hybridization of the O1 and O1' output domains prevents the reaction of Ref-C1-RNA with the O1 DNA Reporter.

#### 5.2. Simulations of the comparator

Simulations of the comparator were conducted using Python's SciPy package (version 1.10.1) using the odeint library. We assumed that both Ref-O1-dART and IFN-O1'-dART without bound protein were transcribed by T7 RNAP with the same rate constant $k_{txn}$ (Figure S19a).

The rate constant of IFN-O1'-RNA binding to Ref-O1-RNA ($k_{th}$) was assumed to be $1 \times 10^6\ M^{-1}s^{-1}$ (Figure S19b)[10]. As in Supplementary Section 2.1, the strand displacement rate constant ($k_{sd}$) for the reaction between Ref-O1-RNA and $[Reporter]_{unreacted}$ was assumed to be $1 \times 10^5\ M^{-1}s^{-1}$ (Figure S19c)[10]. Eqns S6, S7, and S8 show the mass action rate equations that describe the dynamics of the comparator:

$$(S6)\ \frac{d[Ref\_O1\_RNA]}{dt} = k_{txn}[Ref\_O1\_dART] - k_{th}([Ref\_O1\_RNA][IFN\_O1'\_RNA]) \\ -k_{sd}[Ref\_O1\_RNA][Reporter]_{unreacted}$$

$$(S7)\ \frac{d\left[IFN\_O1'\_RNA\right]}{dt} = k_{txn}[IFN\_O1'\_dART]_{unbound} - k_{th}([Ref\_O1\_RNA][IFN\_O1'\_RNA])$$

$$(S8)\ \frac{d[Reporter]_{reacted}}{dt} = k_{sd}[Ref\_O1\_RNA][Reporter]_{unreacted}$$

The binding reaction between IFN-O1'-dART and IFN-γ is assumed to be at thermodynamic equilibrium prior to adding T7 RNAP, at which point the model applies. The concentration of unbound IFN-O1'-dART is determined as described in Supplementary Section 2.2. The $K_{d,apparent}$ of IFN-O1'-dART was assumed to be 8 nM, the value estimated in Supplementary Section 2.3.

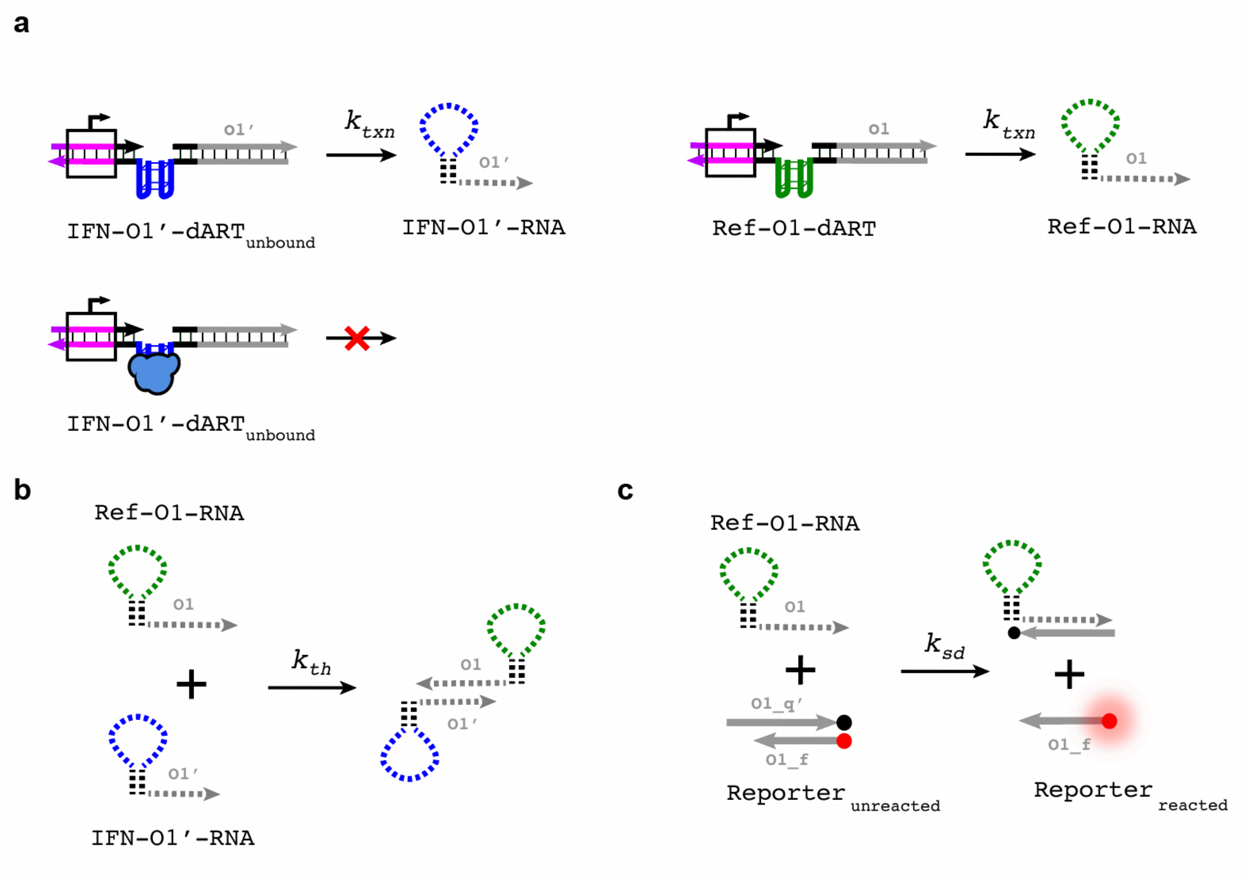


**Figure S19.** The reactions in the model of the comparator. **(a)** IFN-O1'-RNA and Ref-O1-RNA are transcribed from unbound IFN-O1'-dART and Ref-O1-dART, respectively. Transcription reactions are modeled as unimolecular reactions with the same effective reaction rate $k_{txn}$. **(b)** Ref-O1-RNA and IFN-O1'-RNA hybridize *via* a thresholding rate constant $k_{th}$. **(c)** Ref-O1-RNA reacts with the O1 reporter in a 6 base toehold-mediated strand displacement reaction with rate constant $k_{sd}$.

### 5.3. Determining which unbound IFN-O1'-dART concentrations produce high and low levels of comparator output

To design the IFN-γ compartor, we first used simulations (Supplementary Section 5.2) to predict the concentration of unbound IFN-O1'-dART above which little output would be produced from the comparator. To do so, we simulated the comparator with 25 nM of Ref-O1-dART and 250 nM of O1 Reporter and unbound IFN-O1'-dART concentrations ranging from 0 nM to 100 nM. We assumed IFN-O1'-dART would be transcribed at the same rate as Ref-O1-dART.

These simulations predicted that the comparator should produce a relatively low concentration of reacted reporter when the concentration of unbound IFN-O1′-dART is ≥50 nM and should react with all of the reporter within 240 min when the concentration of unbound IFN-O1′-dART is ≤20 nM (Figure S20a). These predictions were confirmed in experiments (Figure S20b-c).

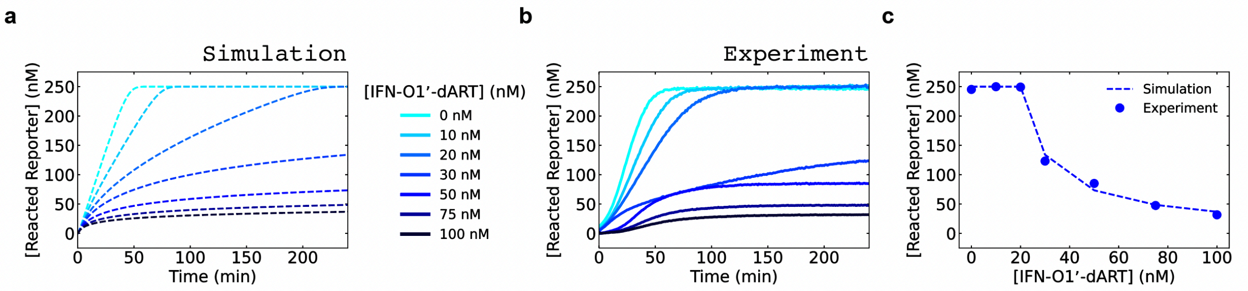


**Figure S20.** Identifying which concentrations of unbound IFN-O1'-dART produce on/off comparator outputs. **(a)** Simulated kinetics of reacted reporter from the comparator with 0 nM to 100 nM of unbound IFN-O1'-dART and 25 nM of Ref-O1-dART ($k_{txn} = 0.004\, s^{-1}$). **(b)** Experimental kinetics of reacted reporter for the comparator with 0 nM to 100 nM of IFN-O1'-dART and 25 nM of Ref-O1-dART. **(c)** Simulated and experimentally measured reacted reporter concentrations after 240 min plotted *vs.* unbound IFN-O1'-dART concentration.

### 5.4. Determining the IFN-γ threshold for digital response of the comparator

Having determined that 50 nM of IFN-O1'-dART would keep reporter signal low in the comparator in Supplementary Section 5.3, we used simulations to identify the concentration of IFN-γ that would repress transcription of IFN-O1'-dART enough to produce a digital ON signal. As described in Section 2.2, we calculated the concentration of unbound IFN-O1'-dART for IFN-γ concentrations of (0, 5, 10, 20, 50, 75, 100, or 200) nM and 50 nM of IFN-O1'-dART (Supplementary Table 5). We then used kinetic simulations of the comparator with these unbound IFN-O1'-dART concentrations to predict the comparator's response for the corresponding input protein concentrations (Figure 5c of the main text). We also used these simulations to predict the behavior of the comparator for different concentrations of Ref-O1-dART (15 nM, 25 nM, 40 nM), which changes the threshold IFN-γ concentration (Figure 5f of the main text).

**Supplementary Table 5.** Predicted unbound dART concentrations were used to produce the simulated reacted reporter kinetics in Figures 5c, 5e, and 5f of the main text. The unbound IFN-O1'-dART concentration for each input protein concentration was calculated as described in Section 2.2 with a $K_{d,apparent}$ of 8 nM, a total IFN-O1'-dART concentration of 50 nM, and IFN-γ concentrations of (0, 5, 10, 20, 30, 50, 75, 100, or 200) nM.

| [IFN-γ] (nM) | [IFN-O1'-dART] (nM) | Rate Parameters | Rate constants |
|---|---|---|---|
| 0 | 50.0 | k_txn | 0.004 $s^{-1}$ |
| 5 | 45.7 | k_th | $1\times10^{6}$ $M^{-1}$ $s^{-1}$ |
| 10 | 41.6 | k_sd | $1\times10^{5}$ $M^{-1}$ $s^{-1}$ |
| 20 | 33.8 | | |
| 30 | 26.9 | | |
| 50 | 16.4 | | |
| 75 | 9.4 | | |
| 100 | 6.2 | | |
| 200 | 2.5 | | |

### 5.5. Dose-response curves of the comparator are similar for different T7 RNAP concentrations

If both IFN-O1'-dART and Ref-O1'-dART are transcribed at similar transcription rates, the dose response of the comparator should not vary significantly with changes in T7 RNAP activity. To test this hypothesis, we repeated the characterization of the IFN comparator in main text Figure 5d using half or twice the T7 RNAP concentration. Similar dose-response curves were observed for 0 nM to 100 nM IFN-γ for the three T7 RNAP concentrations, although as predicted in simulations, the time to saturate the reporter signal decreased with increasing T7 RNAP concentration (Figure S21 and S22).

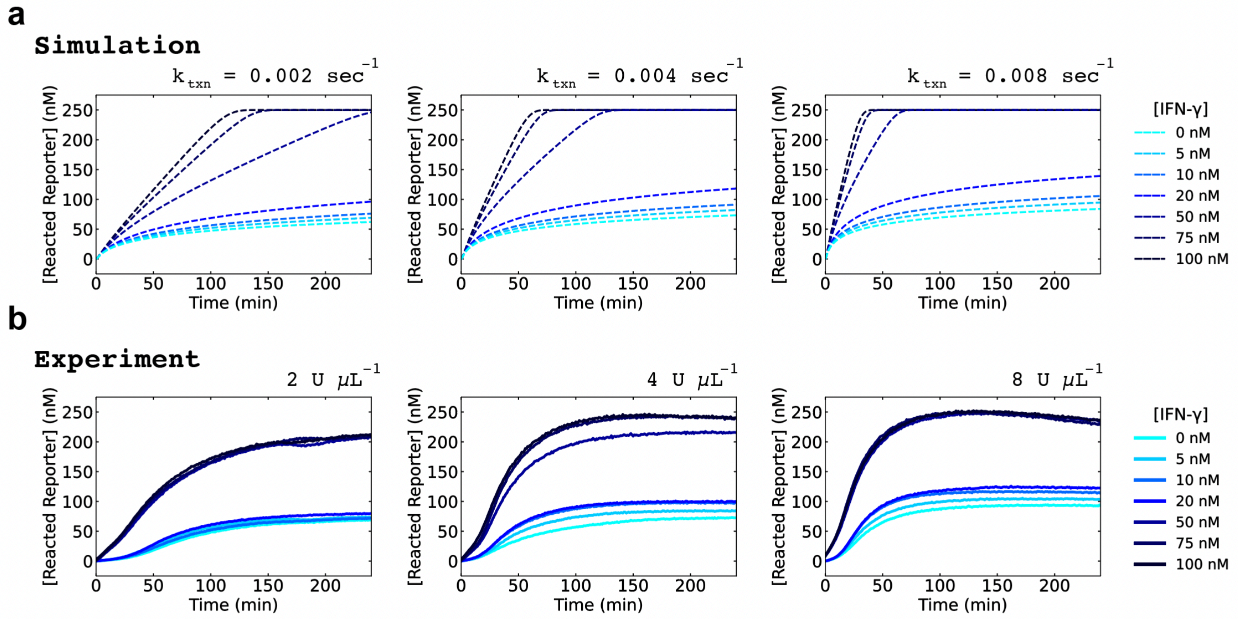


**Figure S21.** Simulated (**a**) and experimental (**b**) reacted reporter kinetics of the comparator with 0 nM to 100 nM of IFN-γ and three different T7 RNAP concentrations. For the simulations, $k_{txn}$ values of 0.002 $s^{-1}$, 0.004 $s^{-1}$, and 0.008 $s^{-1}$ were used to estimate the rates of transcription at T7 RNAP concentrations of 2 U μL$^{-1}$, 4 U μL$^{-1}$, and 8 U μL$^{-1}$, respectively. The amount of T7 RNAP used in each experiment is shown as the title. The 4 U μL$^{-1}$ data is also shown in main text Figure 5d.

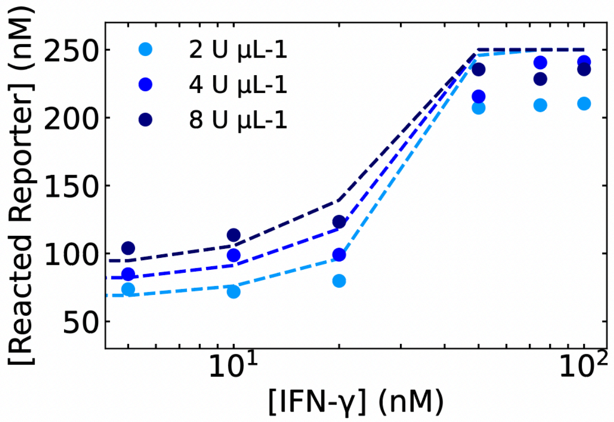


**Figure S22.** Simulated (dashed) and experimental (bold) dose-response curves for the comparator for three different T7 RNAP concentrations as shown in the legend. Reacted reporter concentrations were measured after 240 min of reaction.

### 5.6. Insensitivity of the comparator to potassium ion concentration

The reacted reporter kinetics corresponding to Figure 5h and 5j in the main text are in Figure S23 and S24, respectively.

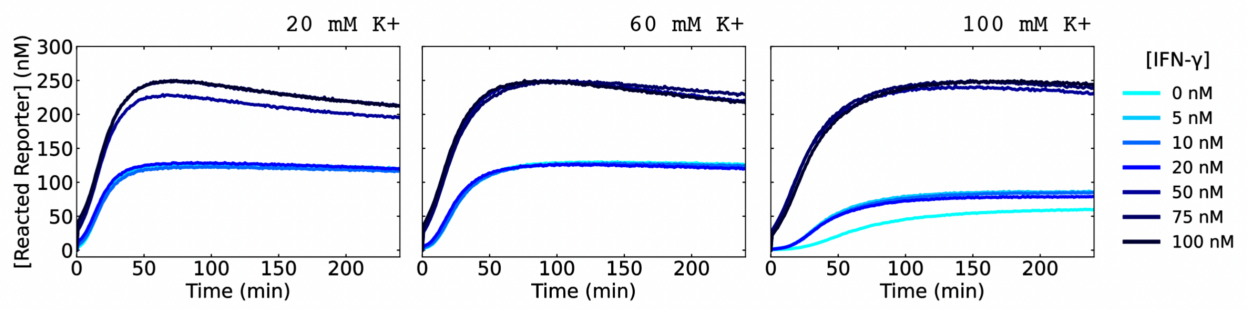


**Figure S23.** Reacted reporter concentrations of the comparator for 0 nM to 100 nM of IFN-γ with varying concentrations of potassium ions (20 mM to 100 mM). The comparator demonstrates digital responses to low (0 nM to 20 nM) and high (50 nM to 100 nM) concentrations of IFN-γ. The dose-response curves in Figure 5h of the main text were obtained from the endpoint reacted reporter concentrations shown in these plots. As [K+] increased, the baseline signal, defined as the concentration of reacted reporter without the input of IFN-γ, decreased. This decrease could be caused by higher [K+] facilitating stable G-quadruplex formation[3], which would reduce the rate of transcription of both dARTs, analogous to reducing the global transcription rate[4].

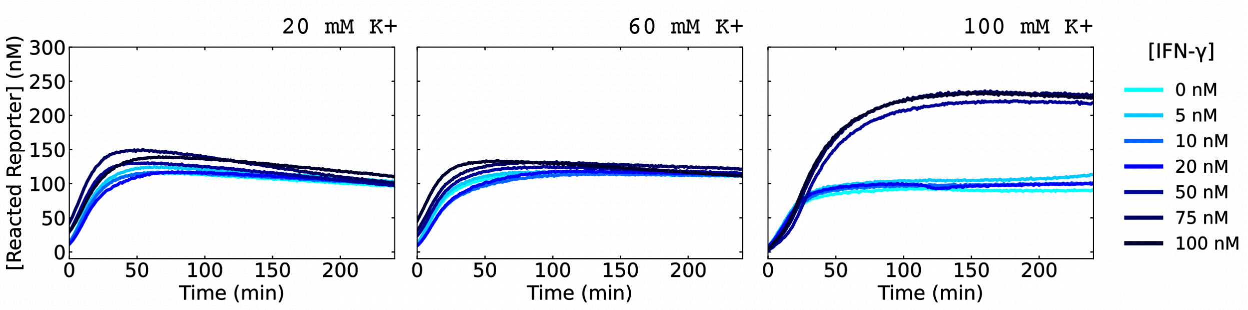


**Figure S24.** Reacted reporter concentrations over time of the dummy comparator, which compares the outputs of 25 nM Dummy-O1-dART and 50 nM IFN-O1'-dART for 0 nM to 100 nM of IFN-γ for three different concentrations of KCl, which are given above the corresponding plots. The dose-response curves in Figure 5j of the main text were obtained from the endpoint reacted reporter concentrations shown in these plots.

### 6. Amplified digital biosensor

The amplified digital biosensor, or amplified comparator, consists of a comparator that compares the outputs of an IFN-γ dART and a reference dART, IFN-C1'-RNA and Ref-C1-RNA, and a blocked genelet that is activated for transcription by Ref-C1-RNA. NUPACK predicts that the C1 and C1' domains of Ref-C1-RNA and IFN-C1'-RNA hybridize as designed (Figure S25). The C1 domain also reacts with a *genelet* (Figure S26), a DNA template whose transcription is regulated by RNA inputs *via* reactions developed previously[1]. Specifically, C1 is the coactivator sequence for the genelet G1O4. G1O4 is in a blocked state because it is bound to the DNA blocker strand B1. The promoter sequence of the resulting complex, G1O4:B1, is incomplete, so it is not transcribed at significant rates. C1 can bind to B1, freeing G1O4. G1O4 can then bind to A1 to produce the complex G1O4:A1 which has a complete promoter. G1O4:A1 can be transcribed to produce O4 RNA that reacts with the reporter.

The amplified comparator also includes the enzyme RNaseH. Ref-C1-RNA bound to B1 can be degraded by RNase H to produce a free B1 strand. Free B1 can bind to G1O4 and can displace A1 from G1O4:A1. Both reactions return the genelet G1O4 to the blocked state G1O4:B1. Free B1 can also bind to intact Ref-C1-RNA. RNase H can also degrade O4-RNA bound to the O4'_q strand of the DNA Reporter, allowing O4'_q and O4_f to hybridize again to reduce fluorescence.

Note that the C1' domain of IFN-C1'-RNA was designed not to be fully complementary to the hairpin in C1. This was done to prevent IFN-C1'-RNA from binding to G1O4, which would block the A1 from binding to G1O4 to activate it.

The 0.02x amplified comparator and 0.02x comparator use the same DNA strands as the amplified comparator and comparator, respectively, each at 0.02 times the concentrations of the amplified comparator and comparator. We compared the responses of these two circuits to IFN-γ concentrations between 0 nM to 4 nM. In its on state, the 0.02x amplified comparator produced up to 250-fold more RNA output than its protein input; the increase in the RNA output between the on state and the baseline state of the comparator was up to 200-fold the input protein concentration (Figure S28a). In contrast, the 0.02x comparator produced negligible reacted O4 reporter for concentration of IFN-γ between 0 nM to 4 nM (Figure S28b).

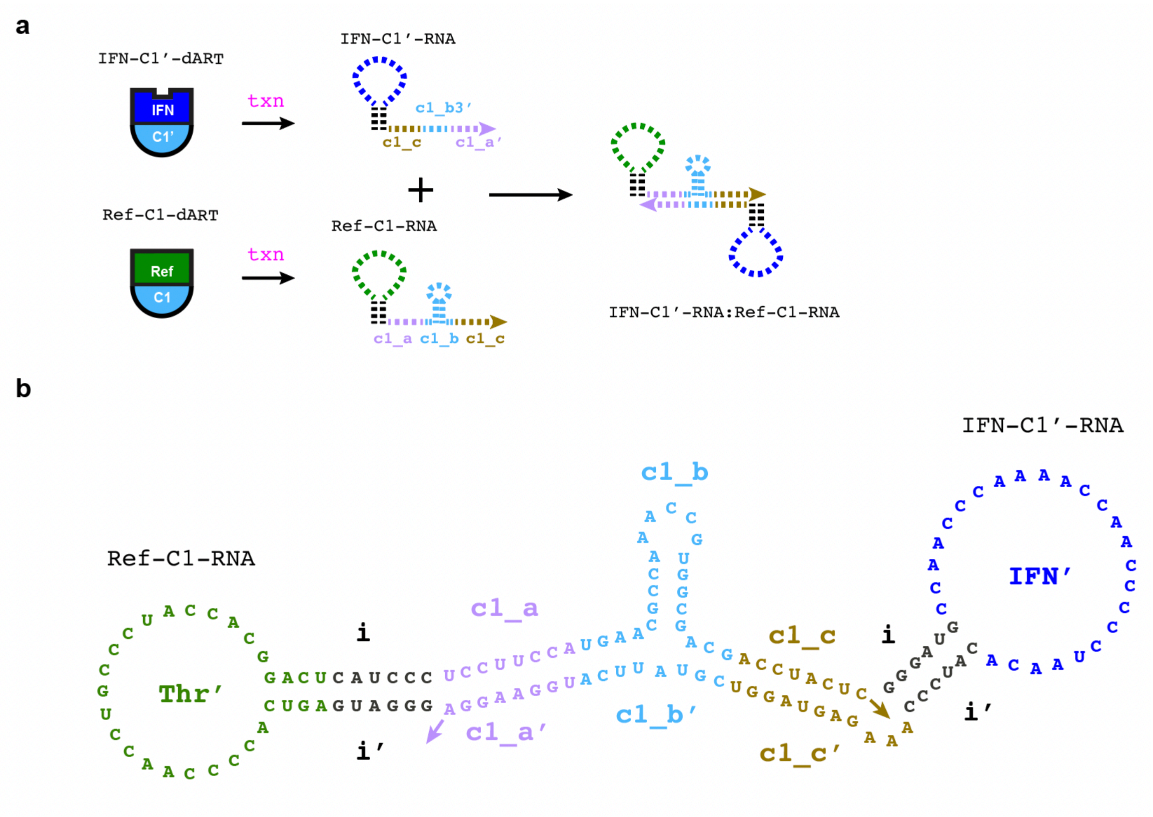


**Figure S25.** Reaction between IFN-C1'-RNA and Ref-C1'-RNA. **(a)** IFN-C1'-RNA is transcribed by unbound IFN-C1'-dART and hybridizes with Ref-C1-RNA. **(b)** The secondary structure of the IFN-C1'-RNA:Ref-C1-RNA complex predicted by NUPACK. The hybridization of the c1 and c1' output domains prevent Ref-C1-RNA from reacting with the G1O4 blocked genelet or B1 (Figure S26) when it is bound to IFN-C1'-RNA.

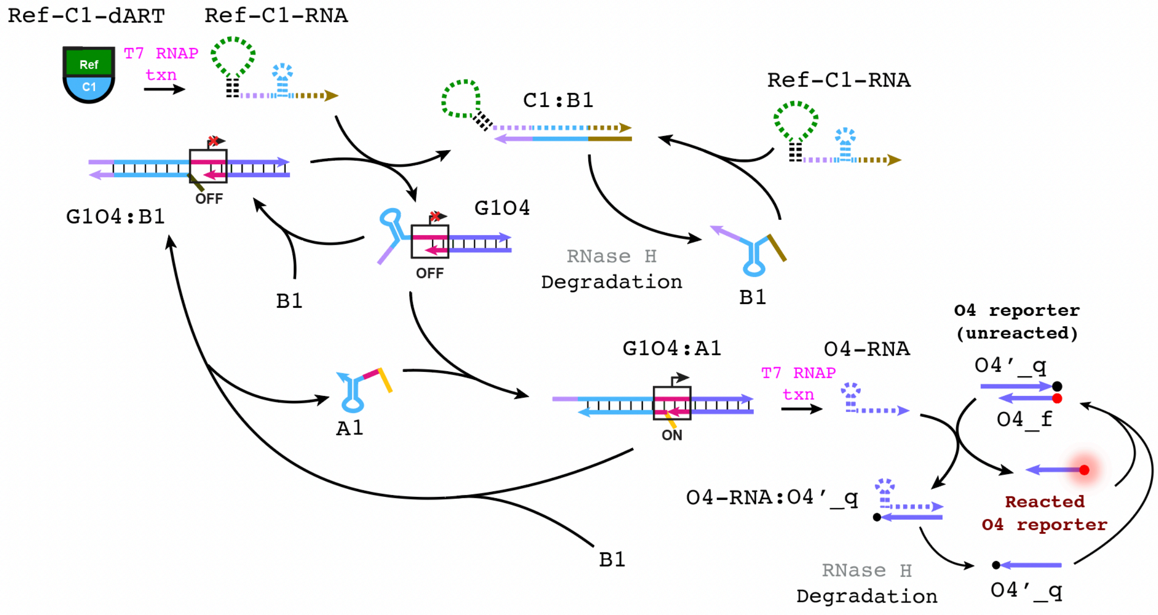


**Figure S26.** Detailed schematic of the genelet reactions in the amplified comparator. When IFN-C1'-dART is bound to its protein ligand, Ref-C1-dART produces Ref-C1-RNA, which displaces B1 from G1O4:B1 to produce G1O4. A1 is then able to bind to G1O4; the resulting complex G1O4:A1 has a complete T7 promoter and can be transcribed to produce O4-RNA. The O4-RNA can react with the O4 DNA Reporter, which is the measured fluorescent output. B1 can bind to G1O4:A1, resulting G1O4:B1 and A1.

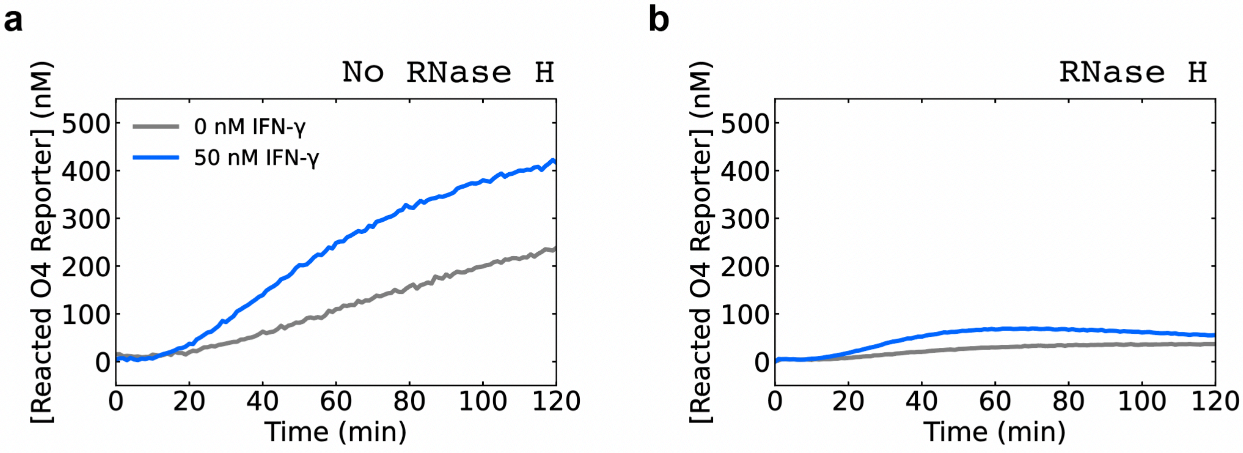


**Figure S27.** Kinetics of the comparator with 0 nM or 50 nM IFN-γ and without **(a)** or with **(b)** RNase H ($4 \times 10^{-3}$ U μL$^{-1}$). The total DNA reporter concentration was 2000 nM.

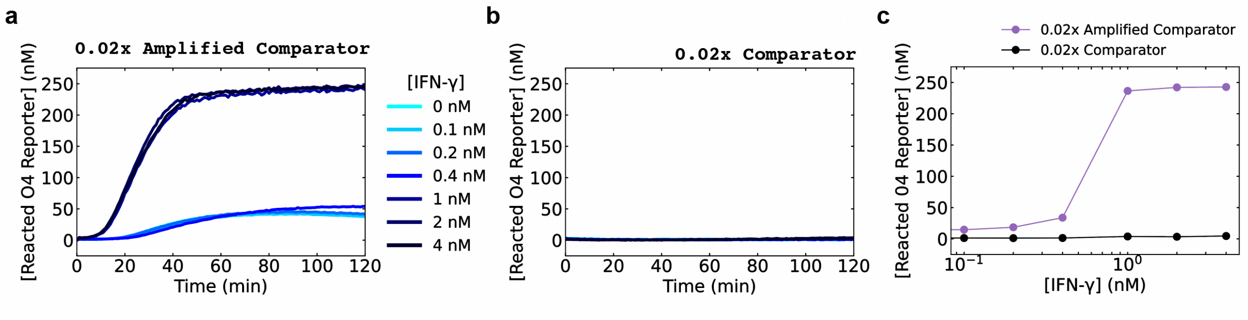


**Figure S28.** Comparing sensitivity of the 0.02x amplified comparator to that of the 0.02x comparator. **(a)** Kinetics of the 0.02x amplified comparator for IFN-γ inputs from 0 nM to 4 nM. (**b**) Kinetics of the 0.02x comparator for IFN-γ inputs from 0 nM to 4 nM. **(c)** Comparison of dose-response curves of the 0.02x amplified comparator and the 0.02x comparator for IFN-γ inputs from 0 nM to 4 nM. For the dose-response curve, the reacted reporter concentrations were measured at 240 min. The kinetic data of the 0.02x amplified comparator is also shown in main text Figure 6d.

### 7. Reagents and their concentrations and experimental protocols

The concentrations listed under Supplementary Tables 6 to 22 are all final concentrations of each component mixed in a total reaction volume of 25 μL per assay.

**Supplementary Table 6.** Measuring reacted report kinetics of IFN-O1-dART when bound to 100 nM of IFN-γ (Figure 2e, main text).

| | Component | Concentration |
|---|---|---|
| 1 | KCl | 0 mM or 100 mM |
| 2 | IFN-O1-dART/Dummy-O1-dART | 10 nM |
| 3 | IFN-γ | 0 nM or 100 nM |
| 4 | O1 DNA Reporter | 100 nM |
| 5 | T7 RNAP | 2 U $\mu L^{-1}$ |
| 6 | YIPP | 1.35 x $10^{-3}$ U $\mu L^{-1}$ |

**Supplementary Table 7.** Dose-response curve for 10 nM of IFN-O1-dART detecting 0 nM to 1000 nM of IFN-γ (Figure 2f and 2g, main text).

| | Component | Concentration |
|---|---|---|
| 1 | IFN-O1-dART | 10 nM |
| 2 | IFN-γ | (0,5,10,25,50,100,250,500,1000) nM |
| 3 | O1 DNA Reporter | 100 nM |
| 4 | T7 RNAP | 2 U $\mu L^{-1}$ |
| 5 | YIPP | 1.35 x $10^{-3}$ U $\mu L^{-1}$ |

**Supplementary Table 8.** Dose-response curve for 1 nM of IFN-O1-dART detecting 0 nM to 100 nM of IFN-γ (Figure 2h and 2i, main text).

| | Component | Concentration |
|---|---|---|
| 1 | IFN-O1-dART | 1 nM |
| 2 | IFN-γ | (0,1,3,5,10,100) nM |
| 3 | O1 DNA Reporter | 100 nM |
| 4 | T7 RNAP | 2 U $\mu L^{-1}$ |
| 5 | YIPP | 1.35 x $10^{-3}$ U $\mu L^{-1}$ |

**Supplementary Table 9.** Measuring the dose-response curves for different dARTs (Figure 3b, main text).

| | Component | Concentration |
|---|---|---|
| 1 | IFN-O1-dART/Thr-O1-dART/IL6-O1-dART/TNF-O1-dART | 10 nM |
| 2 | IFN-γ/Thrombin/IL-6/ TNF-α | (0,5,10,100,1000) nM |
| 3 | O1 DNA Reporter | 100 nM |
| 4 | T7 RNAP | 2 U $\mu L^{-1}$ |
| 5 | YIPP | 1.35 x $10^{-3}$ U $\mu L^{-1}$ |

**Supplementary Table 10.** Selectivity of dARTs (Figure 3c, main text; Figure S11). For each assay, one of the dARTs and O1 DNA Reporter were mixed in ARTIST reaction condition, at final concentrations of 10 nM and 100 nM, respectively. 100 nM of BSA, IFN-γ, thrombin, IL-6, or TNF-α was also included into the assay.

| | Component | Concentration |
|---|---|---|
| 1 | IFN-O1-dART/Thr-O1-dART/IL6-O1-dART/TNF-O1-dART/Dummy-O1-dART | 10 nM |
| 2 | None/BSA/IFN-γ/Thrombin/IL-6/ TNF-α | 100 nM |
| 3 | O1 DNA Reporter | 100 nM |
| 4 | T7 RNAP | 2 U $\mu L^{-1}$ |
| 5 | YIPP | 1.35 x $10^{-3}$ U $\mu L^{-1}$ |

**Supplementary Table 11.** Modularity of dART output (Figure 3e, main text).

| | Component | Concentration |
|---|---|---|
| 1 | IFN-O1-dART/IFN-O2-dART/IFN-O3-dART | 10 nM |
| 2 | IFN-γ | (0,5,10,100,1000) nM |
| 3 | O1 DNA Reporter | 100 nM |
| 4 | T7 RNAP | 2 U $\mu L^{-1}$ |
| 5 | YIPP | 1.35 x $10^{-3}$ U $\mu L^{-1}$ |

**Supplementary Table 12.** Evaluation of efficient T7 RNAP transcription of KMy-O1-dART without/with 400 μM of kanamycin (Figure S16).

| | Component | Concentration |
|---|---|---|
| 1 | KMy-O1-dART | 10 nM |
| 2 | Kanamycin | 0 μM or 400 μM |
| 3 | O1 DNA Reporter | 100 nM |
| 4 | T7 RNAP | 2 U $\mu L^{-1}$ |
| 5 | YIPP | 1.35 x $10^{-3}$ U $\mu L^{-1}$ |

**Supplementary Table 13.** Steady-state analog output signals of dARTs with varying ligand concentrations (Figure 4b and 4c main text; Figure S17).

| | Component | Concentration |
|---|---|---|
| 1 | IFN-O1-dART/Thr-O1-dART/IL6-O1-dART/TNF-O1-dART | 10 nM |
| 2 | IFN-γ/Thrombin/IL-6/ TNF-α | (0,5,10,100,1000) nM |
| 3 | O1 DNA Reporter | 100 nM |
| 4 | T7 RNAP | 2 U/μL |
| 5 | YIPP | 1.35 x $10^{-3}$ U/μL |
| 6 | RNase H | 2.00 x $10^{-3}$ U/μL |

**Supplementary Table 14.** Experimental setup to determine threshold concentration of IFN-O1'-dART for the IFN-γ comparator (Figure S20).

| | Component | Concentration |
|---|---|---|
| 1 | IFN-O1'-dART | (0,10,20,30,50,75,100) nM |
| 2 | Ref-O1-dART (Thr-O1-dART) | 25 nM |
| 3 | O1 DNA Reporter | 250 nM |
| 4 | T7 RNAP | 4 U $\mu L^{-1}$ |
| 5 | YIPP | 1.35 x $10^{-3}$ U $\mu L^{-1}$ |

**Supplementary Table 15.** Measuring response of the IFN-γ comparator with varying IFN-γ concentrations (Figure 5d and 5e, main text).

| | Component | Concentration |
|---|---|---|
| 1 | IFN-O1′-dART | 50 nM |
| 2 | Ref-O1-dART (Thr-O1-dART) | 25 nM |
| 3 | IFN-γ | (0,5,10,20,50,75,100) nM |
| 4 | O1 DNA Reporter | 250 nM |
| 5 | T7 RNAP | 4 U $\mu L^{-1}$ |
| 6 | YIPP | 1.35 x $10^{-3}$ U $\mu L^{-1}$ |

**Supplementary Table 16.** Shifting the dose-response curves of the comparator (Figure 5f, main text).

| | Component | Concentration |
|---|---|---|
| 1 | IFN-O1′-dART | 50 nM |
| 2 | Ref-O1-dART (Thr-O1-dART) | (15,25,40) nM |
| 3 | IFN-γ | (0,5,10,30,50,100,200) nM |
| 4 | O1 Reporter | 250 nM |
| 5 | T7 RNAP | 4 U $\mu L^{-1}$ |
| 6 | YIPP | 1.35 x $10^{-3}$ U $\mu L^{-1}$ |

**Supplementary Table 17.** Dose-response curves of the comparator with varying T7 RNAP concentrations (Figure S21 and S22).

| | Component | Concentration |
|---|---|---|
| 1 | IFN-O1′-dART | 50 nM |
| 2 | Ref-O1-dART (Thr-O1-dART) | 25 nM |
| 3 | IFN-γ | (0,5,10,20,50,75,100) nM |
| 4 | O1 Reporter | 250 nM |
| 5 | T7 RNAP | 2/4/8 U $\mu L^{-1}$ |
| 6 | YIPP | 1.35 x $10^{-3}$ U $\mu L^{-1}$ |

**Supplementary Table 18.** Testing K+ sensitivity of the comparator with Ref or Dummy dART (Figure 5h and 5j, main text; Figure S23 and S24).

| | Component | Concentration |
|---|---|---|
| 1 | KCl | (20,60,100) mM |
| 2 | IFN-O1′-dART | 50 nM |
| 3 | Ref-O1-dART or Dummy-O1-dART | 25 nM |
| 4 | IFN-γ | (0,5,10,20,50,75,100) nM |
| 5 | O1 DNA Reporter | 250 nM |
| 6 | T7 RNAP | 4 U $\mu L^{-1}$ |
| 7 | YIPP | 1.35 x $10^{-3}$ U $\mu L^{-1}$ |

**Supplementary Table 19.** The amplified comparator (Figure 6c, main text).

| | Component | Concentration |
|---|---|---|
| 1 | IFN-C1′-dART | 50 nM |
| 2 | Ref-C1-dART (Thr-C1-dART) | 25 nM |
| 3 | G1O4:B1 | 100 nM |
| 4 | A1 | 200 nM |
| 5 | IFN-γ | 0 nM or 50 nM |
| 6 | O4 DNA Reporter | 2000 nM |
| 7 | T7 RNAP | 4 U $\mu L^{-1}$ |
| 8 | YIPP | 1.35 x $10^{-3}$ U $\mu L^{-1}$ |
| 9 | RNase H | 4.00 x $10^{-3}$ U $\mu L^{-1}$ |

**Supplementary Table 20.** Unamplified comparator (Figure S27; RNase H was added to Figure S27b).

| | Component | Concentration |
|---|---|---|
| 1 | IFN-O4′-dART | 50 nM |
| 2 | Ref-O4-dART (Thr-O4-dART) | 25 nM |
| 3 | IFN-γ | 0 nM or 50 nM |
| 4 | O4 DNA Reporter | 2000 nM |
| 5 | T7 RNAP | 4 U $\mu L^{-1}$ |
| 6 | YIPP | 1.35 x $10^{-3}$ U $\mu L^{-1}$ |
| 7 | RNase H | 4.00 x $10^{-3}$ U $\mu L^{-1}$ |

**Supplementary Table 21.** Dose-response curve of the 0.02x amplified comparator (Figure 6d and 6e, main text; Figure S28a and S28c).

| | Component | Concentration |
|---|---|---|
| 1 | IFN-C1′-dART | 1 nM |
| 2 | Ref-C1-dART (Thr-C1-dART) | 0.5 nM |
| 2 | G1O4:B1 | 20 nM |
| 3 | A1 | 100 nM |
| 5 | IFN-γ | (0,0.1,0.2,0.4,1,2,4) nM |
| 6 | O4 Reporter | 250 nM |
| 7 | T7 RNAP | 4 U $\mu L^{-1}$ |
| 5 | YIPP | 1.35 x $10^{-3}$ U $\mu L^{-1}$ |
| 6 | RNase H | 4.00 x $10^{-3}$ U $\mu L^{-1}$ |

**Supplementary Table 22.** Dose-response curve of the 0.02x comparator to detect 0 nM to 4 nM of IFN-γ (Figure S28b and S28c).

| | Component | Concentration |
|---|---|---|
| 1 | IFN-O4'-dART | 1 nM |
| 2 | Ref-O4-dART (Thr-O4-dART) | 0.5 nM |
| 3 | IFN-γ | (0,0.1,0.2,0.4,1,2,4) nM |
| 4 | O4 Reporter | 250 nM |
| 5 | T7 RNAP | 4 U $\mu L^{-1}$ |
| 6 | YIPP | 1.35 x $10^{-3}$ U $\mu L^{-1}$ |
| 7 | RNase H | 4.00 x $10^{-3}$ U $\mu L^{-1}$ |
